# Engineered biological neural networks as basic logic operators

**DOI:** 10.1101/2024.12.23.630065

**Authors:** Joël Küchler, Katarina Vulić, Haotian Yao, Christian Valmaggia, Stephan J. Ihle, Sean Weaver, János Vörös

## Abstract

We present an *in vitro* neuronal network with controlled topology capable of performing basic Boolean computations, such as NAND and OR. Neurons cultured within polydimethylsiloxane (PDMS) microstructures on high-density microelectrode arrays (HD-MEAs) enable precise interaction through extracellular voltage stimulation and spiking activity recording. The architecture of our system allows for creating non-linear functions with two inputs and one output. Additionally, we analyze various encoding schemes, comparing the limitations of rate coding with the potential advantages of spike-timing-based coding strategies. This work contributes to the advancement of hybrid intelligence and biocomputing by offering insights into neural information encoding and decoding with the potential to create fully biological computational systems.

## 1 Introduction

Information processing and learning in artificial intelligence (AI) systems, while diverse, is well-known and understood. At its core, AI learning involves training algorithms that iteratively adjust internal parameters by minimizing the difference between predicted and actual outputs through techniques like backpropagation and gradient descent. Among other aspects, the learning process in AI systems typically involves the use of various activation functions, such as sigmoid, ReLU, or softmax, which determine the output of each neuron based on its inputs [1]. These activation functions are crucial in enabling the system to learn complex patterns and relationships within the data [2, 3, 4]. One of the key aspects of information processing in AI systems is the use of artificial neural networks (ANNs), which are inspired by the structure and function of the human brain [1].

On the contrary, the mechanisms by which the human brain encodes, processes, and propagates information are complex and not well understood. It appears that different regions of the brain are specialized for various functions, each exhibiting unique properties in terms of information encoding, processing precision, and memory duration [5, 6, 7]. Humans learn quickly and efficiently. They are able to generalize and thus still outperform AI systems [8]. To accomplish efficient information transmission (and its storage), neurons in the brain communicate through synapses, which transmit signals from one neuron to another. The process involves the release of neurotransmitters, which cross the synaptic cleft and bind to receptors on the receiving neuron. This can initiate an action potential that propagates along the neuron and triggers the release of neurotransmitters at the next synapse. The biological processes that result in the creation of an action potential (output) upon receiving an input can be described as biological activation functions [9]. This chain of events allows for the transmission of information across different parts of the brain.

In recent years, there has been an emphasis on leveraging biological mechanisms to improve artificial learning, mainly in efficiency and generalizability. These include the development of neuromorphic hardware [10, 11, 12] and the implementation of spiking neural networks (SNNs) [13], which are inspired by biological neural networks [14]. Neuromorphic computing replicates the structure of the human brain and may enable highly efficient computing through parallel computation and adaptive learning [15]. SNNs offer an energy-efficient alternative to traditional ANNs due to their spike-based communication and computation mechanisms [16, 17]. The main limitation of SNNs and neuromorphic hardware is that, as mentioned before, the biological mechanisms of learning and information processing are not fully understood, which complicates accurate implementation [18, 19]. Important questions relate to identifying the aforementioned biological activation functions and finding proper input/output (I/O) encoding mechanisms.

One perspective on biological behavior in terms of information encoding, transmission, and decoding postulates that neurons operate similarly to communication channels [20]. In this view, neurons receive inputs, process information, and then produce, to some extent, noisy outputs, effectively acting as conduits for transmitting and transforming information within the brain. This theory draws on the analogy to information theory, where communication channels transmit data between a sender and a receiver [21, 22, 23].

Several coding theories intend to explain how neurons represent and communicate information. Key frameworks include rate coding and temporal coding, such as phase coding, inter-spike interval (ISI) coding, and time-to-first spike (TTFS) coding. Rate coding represents information by the frequency of spikes within a specific time frame. Research suggests that the cortex mainly uses rate coding, as it is resistant to noise due to the averaging effect over time [24]. This makes rate coding particularly effective for reliable information transfer, even in highly variable environments. Rate coding carries an implication of temporal averaging which can work well when the stimulus is constant, slowly varying, or does not require a fast reaction. Outside these assumptions, it can be limiting [25].

Phase coding uses the timing of spikes relative to a periodic signal, which helps capture temporal patterns in sensory information [26]. This approach has been shown to strengthen the stability of information carried by spatial and temporal spike patterns, improving the accuracy of sensory representations [26].

Similarly, TTFS coding, which focuses only on the timing of the first spike in response to a stimulus, is valued for its speed in transmitting information, especially in fast sensory environments [27]. Precise spike timing in TTFS coding plays a key role in interpreting sensory signals [27, 28].

ISIs are important in neural coding, as they provide information on the statistical properties of stimuli. Neurons can adapt to different signal features by adjusting their firing intervals [29]. ISIs can reflect stimulus variability, highlighting the role of spike timing patterns in sensory processing [29]. Analyzing ISI variability also provides insights into neural behavior and the characteristics of incoming input signals [30]. This is particularly important in cases where changes in neural behavior need to be quantified, such as during neuroplasticity, *i*.*e*., the learning processes of neurons.

The encoding mechanisms and activation processes of biological neurons must be further explored in neural systems, particularly in the context of developing biologically inspired AI systems. While numerous *in silico* studies have explored I/O transformations across various complexity levels of neural systems [31, 32, 33, 34, 35], studying and observing these phenomena in living biological systems remain considerably more limited and challenging [36, 37, 38]. Since currently the brain as a whole represents an incomprehensible complex system, it is helpful, if not essential to break down the complexity to understand the processes of communication and information transmission. One way to obtain well-defined and reproducible biological neural systems suitable for studying such phenomena is to form engineered *in vitro* neuronal networks with controlled topology and defined directionality and connectivity [39, 40, 41]. Additionally, it is necessary to interact with the system in order to provide controlled inputs and to read out the corresponding outputs. One way to achieve network directionality and feed-forward information flow is to culture neurons inside custom-designed polydimethylsiloxane (PDMS) microstructures and to place them on top of (high-density) microelectrode arrays (HD-MEAs). This allows for studying I/O transformations of simplified neural systems by allowing both the observation of the system through the extracellular recording and the manipulation of the system through extracellular voltage stimulation [42, 43, 44, 45, 38]. This methodology has recently been applied to leverage biological systems for various forms of problem-solving [37, 38, 46].

In this work, we present a feedforward biological neural network with a two-input-one-output topology. The topology is controlled by confining neuronal growth within PDMS microstructures. We interact with a network through high density microelectrode arrays. We encode external inputs using specific stimulation patterns and represent output information through different encoding schemes. First, we explore the limitations of information transmission in our system through simple encoding schemes. We identify the optimal encoding strategy and valorize the network’s reproducible behavior to create non-linear biological transfer functions and perform fundamental logic operations. Characterizing these concepts within biological systems could provide valuable insights into the validity and relevance of the principles currently applied in bio-inspired computing.

## 2 Materials and Methods

### 2.1 PDMS microstructures

PDMS microstructures for cell and axon guidance were designed using CAD software (AutoCAD 2021) and fabricated by Wunderlichips (Zurich, Switzerland). The fabrication process is elaborated in previous publications [47, 48]. In this study, all microstructures feature wells for cell seeding, microchannels impermeable to soma but accessible to neurites, and submicron tunnels designed to prevent axonal outgrowth while allowing penetration of dendritic spines to assure the directionality of action potential propagation. Schematics illustrating these microstructures are presented in Fig. 1. The full thickness of the PDMS microstructure measures 75 *μ*m, with 2 *μ*m high microchannels, and a submicron tunnel area with a height of 600 nm. The microstructure was designed to engineer feed-forward neuronal networks consisting of two presynaptic (input) nodes connecting to the postsynaptic (output) node. These networks will from now on be referred to as 2-in-1-out networks. Directionality is achieved by using 250 nm-wide PDMS tunnels (see Fig. 1Biii) that restrict the input axon from growing through, while still allowing dendritic spines from output neurons to penetrate and form synaptic connections with the input (for more details, see previous work [47].

**Fig. 1:**
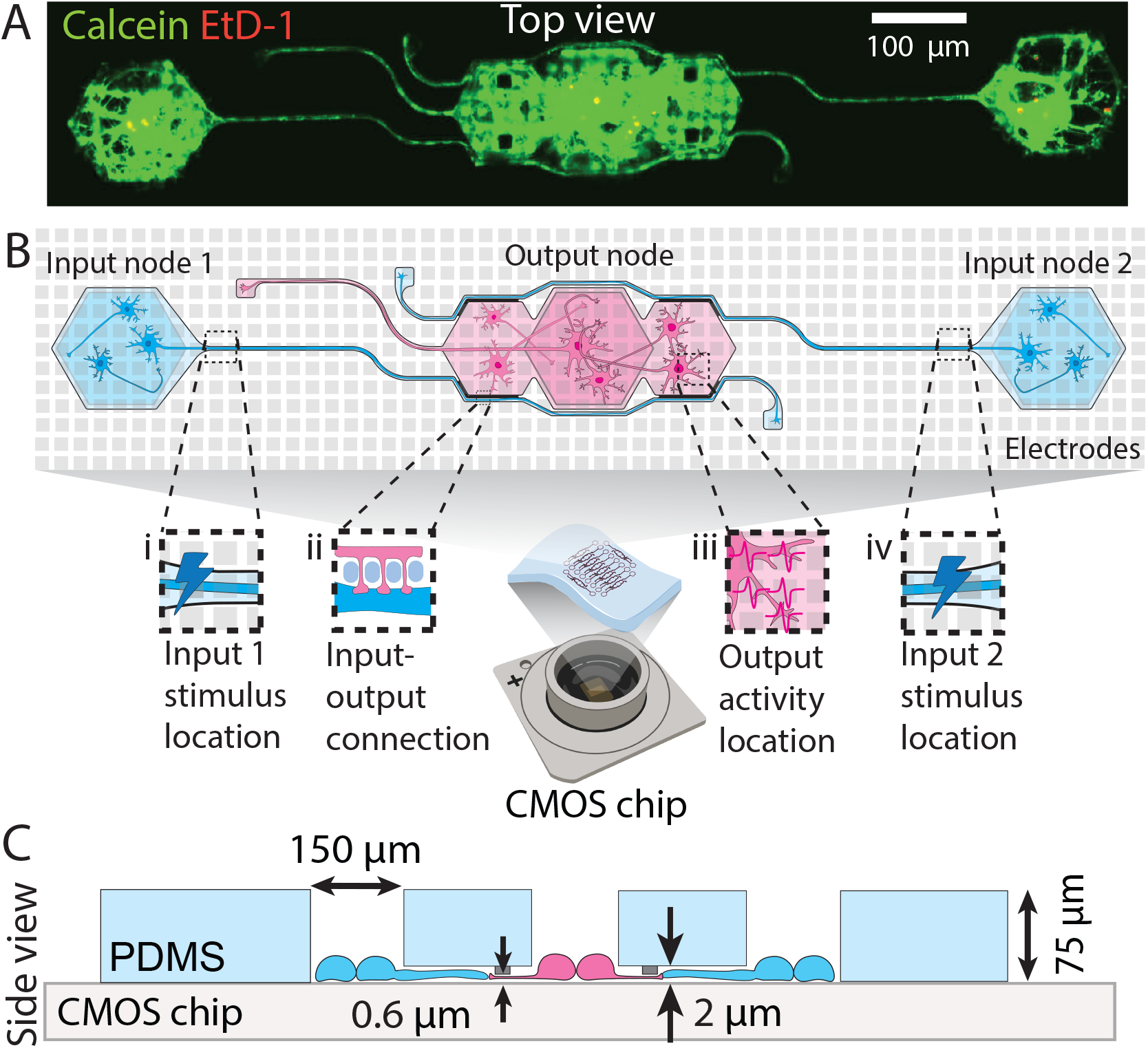
Engineered feedforward neuronal networks on a chip. **(A)** Microscopy example image of a neuronal network defined by a PDMS microstructure. The network consists of two input nodes with microchannels guiding the input axons toward the output node. **(B)** Top view schematic of a network describing (i) the location of the external stimulus applied at the beginning of the first input axon channel, (ii) the putative synaptic connection formed between the input axon and output dendrites through submicron tunnels, (iii) part of the area where output action potentials are recorded from, and (iv) the location of the external stimulus applied at the beginning of the second input axon channel. **(C)** Cross-section schematic of the same microstructure showing the microchannel and submicron tunnel heights, well opening sizes, and the total PDMS thickness.

### 2.2 Microelectrode arrays

Microelectrode arrays (MEAs) used in the experiments are high-density complementary metal-oxide-semiconductor (HD-CMOS) MEAs. The small MaxOne HD-MEA chips with flat surface topology (Maxwell Biosystems, Switzerland) consist of 26 400 electrodes distributed on a 3.85 × 2.10 mm^2^ surface with an electrode pitch of 17.5 *μ*m. It is possible to record from 1020 electrodes simultaneously. These electrodes can be chosen arbitrarily and are routed to the amplifiers through a switch matrix. The data is sampled at 20 kHz. Extracellular stimulation is possible on the same electrodes and is performed with 32 independent stimulation buffers.

### 2.3 Substrate preparation

The preparation begins with the pre-cleaning of the reused MEA, which can be omitted if the new sterile chip is used. Cleaning involves leaving a chip in 4 % Tergazyme (1304-1, Alconox) overnight, rinsing with ultrapure water (Milli-Q, Merck-MilliPore), and then leaving the chips for circa 30 min in 70% ethanol solution. This is followed by three rinses with ultrapure water. Subsequently, the MEA surface is dried as much as possible using a nitrogen gun. Next, the chip is functionalized with Poly-D-Lysine (PDL) (P6407, Sigma Aldrich). The solution is 0.1 mg/mL of PDL in phosphate-buffered saline (PBS) (10010-023, ThermoFisher). 100 *μ*L of the PDL solution is pipetted onto the chip surface, and the chip is left for a minimum of 45 minutes before the chip is rinsed three times with Milli-Q water to remove unbound PDL polymers. The chip is then dried with a nitrogen gun. PDMS microstructure is then placed on the chip surface with tweezers, ensuring that it is as straight as possible and avoiding contact of the tweezers with the chip surface. Once positioned, the microstructure is gently pressed flat onto the chip. Subsequently, 1 mL of warm PBS is added, and the chip is desiccated for a minimum of 5 minutes or until no bubbles are observed on the surface of the microstructure.

### 2.4 Visualization of PDMS microstructure with a CMOS MEA

We employ a custom Python script that generates an voltage map showing the location of electrodes beneath the PDMS microstructure. The example of an voltage map for this PDMS microstructure can be seen in Fig. S1. This enables us to locate and select the electrodes covering individual networks within the PDMS microstructure. The method is described in detail in our previous publication [44].

### 2.5 Cell culture

The cells were cultured in NeuroBasal medium (NB) (21203-049, ThermoFisher) [49]. Fresh NB complete medium was prepared, consisting of a 2% solution of B-27 supplement (17504-044), a 1% solution of Penicillin-Streptomycin (P-S) (15070-063), and a 1% solution of GlutaMAX (35050-061), all sourced from ThermoFisher.

#### 2.5.1 Cell dissociation

Primary hippocampal neurons were obtained from E18 embryos of pregnant Sprague-Dawley rats (EPIC, ETH Phenomics Center) for use in the experiments. All animal procedures were approved by the Cantonal Veterinary Office Zurich. The embryonic neuronal tissue was dissected and stored in Hibernate E medium (Thermo PN) on ice. Cell dissociation commenced by digesting the tissue in a solution containing 50 mg/mL Bovine Serum Albumin (BSA) (A7906, Sigma-Aldrich), 1.8 mg/mL D-glucose (Y0001745, Sigma-Aldrich), and 0.5 mg/mL papain (P5306, Sigma-Aldrich) dissolved in sterile PBS. Prior to dissociation, the solution was warmed to 37°C, filtered through a 0.2 *μ*m filter, and supplemented with 1 mg/mL DNAse (D5025, Sigma-Aldrich). The tissue was incubated in the papain solution for 15 minutes at 37°C, followed by replacement with NB medium containing 10% fetal bovine serum (10500056, ThermoFisher) to halt the digestion process. Subsequently, two washes with NB medium were performed, with a 5-minute interval between each wash. Trituration was then carried out, followed by cell counting using the Cell Countess system (Invitrogen). Cells dissociated from a single pregnant rat were considered as one biological replicate.

#### 2.5.2 Cell seeding and maintenance

Prior to cell seeding onto the chip substrate, PBS was gradually replaced with NB complete medium, avoiding the reformation of air bubbles in the microchannels. Cells were seeded in suspension with a concentration of 70,000 cells per MEA (which corresponds to roughly 250 cells/mm^2^). MEA chips were then transferred into the incubator at 37 °C, 90% humidity and 5% CO_2_. 20-30 min upon seeding, cells were re-suspended in the chip by pipetting up and down to increase the probability of more cells falling into the PDMS wells. The first medium exchange was done one day after seeding to remove the dead cells and cell debris and the medium was then exchanged twice per week.

### 2.6 Experimental Setup

The experimental setup involved stimulation and recording of the neurons topologically constrained to form 2 input node and 1 output node networks (see Fig. 1A). Each chip contained up to 10 independent networks The experiments were performed in the third week of culture to ensure the functional maturity of the network [50].

#### 2.6.1 Network selection

Networks were selected based on the activity in each node. Up to three 2-in-1-out networks were tested simultaneously due to restrictions on the number of recording channels (approx. 280 electrodes per network, up to 1020 recording channels). Only networks that showed activity in three nodes and the connecting channel were used for the analysis. The stimulation electrode location was chosen based on previous research [51] and will be further discussed in Section 3.2.

#### 2.6.2 Data collection

Data was collected on day *in vitro* (DIV) 21 or 22. The MaxOne headstage with the chip was placed in a custom-designed incubator at 36 °C, 90% humidity, and 5% CO_2_ [52]. The temperature increase due to the heat up of the recording headstage was compensated by setting the temperature lower than 37°C. Multiple recording headstages allowed for recording from up to four MEA chips simultaneously, meaning that on each experiment trial, up to 12 networks could be recorded. In each experiment trial, multiple stimulation paradigms were tested, resulting in continuous data collection over multiple hours.

#### 2.6.3 Stimulation and recording paradigms

Two types of experiments were performed to explore the response of the network to external stimuli: voltage amplitude-modulated experiments and frequency-modulated experiments. In both setups, biphasic pulses with a duration of 400 *μ*s were delivered periodically to the MEA. For the amplitude-modulated experiments, the stimulation frequency was fixed at 4 Hz, as established previously [45]. The stimulation amplitude ranged from 0 mV to a maximum of 800 mV [51]. In the frequency-modulated experiments, the inter-stimulus time was altered while the pulse amplitude was kept constant at 400 mV. The stimulation frequency ranged from 1 Hz to 80 Hz. Each sequence of same-amplitude or same-frequency stimuli will be further referred to as a pulse set, and each full amplitude or frequency modulation experiment will be referred to as a stimulation set. The recording of the neuronal activity was conducted simultaneously during stimulation. Output signals were recorded on electrodes in the output node (see Fig. 1Biv) and Fig. 2A). A time window of up to 10 ms after the input stimulus was considered as the output. For a modulation experiment, each pulse set was recorded once for a minimum of 30 seconds. The order of pulse sets was randomized. Further, it was ensured that the number of iterations used for the analysis was equivalent across all parameters within each specific experimental condition.

**Fig. 2:**
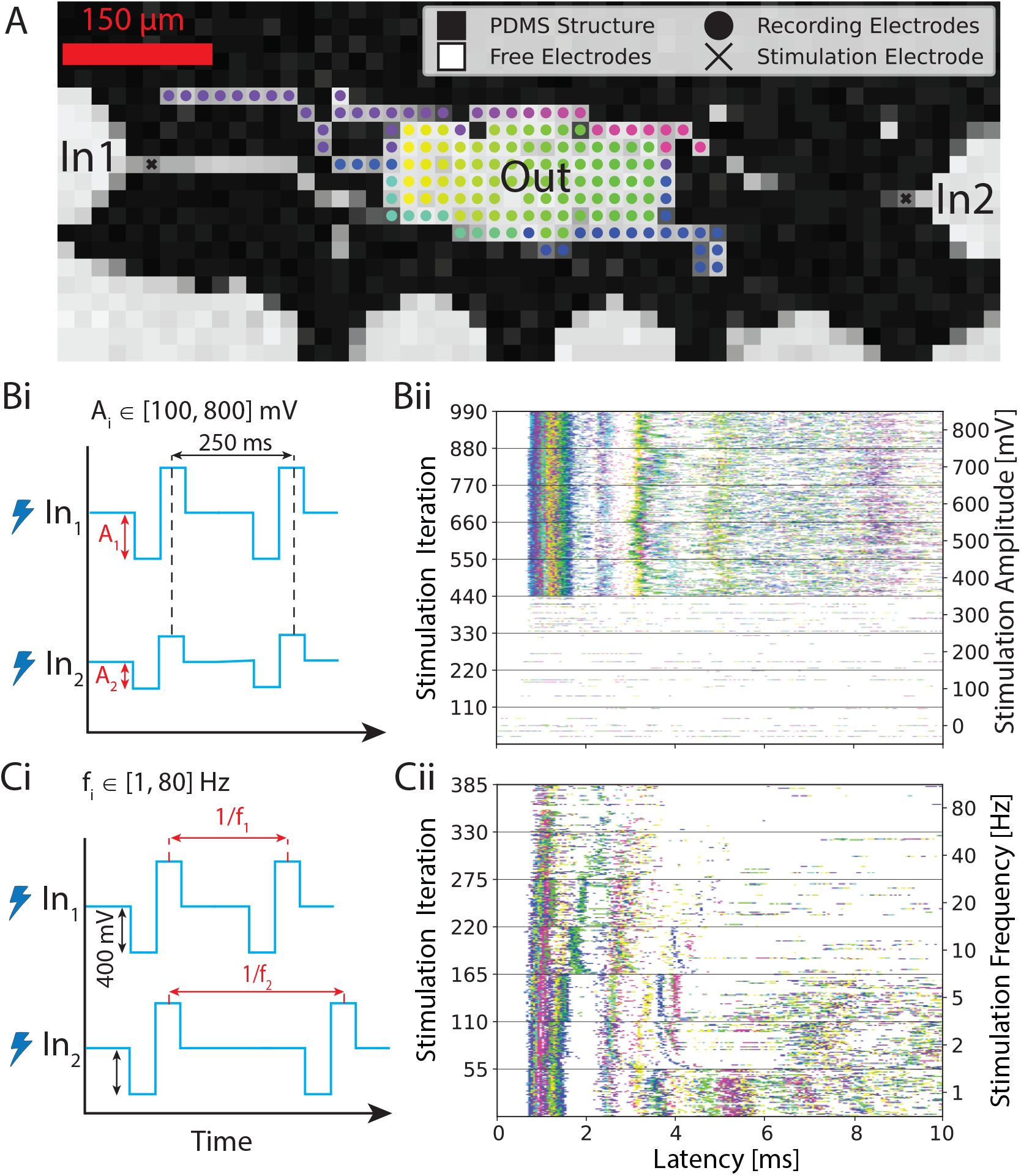
Overview of the experiment. **(A)** Voltage map showing the PDMS microstructure on a CMOS MEA. We stimulate inputs inside the axon channels and record the response from the electrodes at the output. We color-coded each recording electrode. **(Bi)** First input variant: sweeping the input voltage amplitude from 100 to 800 mV with 100 mV step size while keeping the stimulation frequency constant at 4 Hz. **(Bii)** Corresponding post-stimulus raster plots. In this case, the same stimulus is simultaneously applied 110 times for each amplitude iteration through both input electrodes. The first 10 ms of the output response is plotted after each stimulus. **(Ci)** Second input variant: sweeping through stimulation frequency from 1 to 80 Hz while keeping the stimulus voltage amplitude constant at 400 mV. **(Cii)** Corresponding post-stimulus raster plot. In this case, stimulus at the same frequency is simultaneously applied at both input electrode 1 and input electrode 2. The output response is plotted 10 ms after each stimulus.

#### 2.6.4 Data Analysis

Data was processed using a custom-designed Python processing pipeline. The stimulation artifact of each recording electrode was cut out and linearly interpolated (see Section S3 for a detailed explanation). The signal was then filtered using a second-order Butterworth high-pass filter with a cutoff frequency of 200 Hz. It was further smoothed with a 2nd-order Savitzky-Golay filter [53] using a window size of 0.25 ms to remove high-frequency fluctuations irrelevant for spike detection. Spike times were then determined using a smoothed nonlinear energy operator (SNEO) with lag *k* = 3 [54]. Peaks exceeding 20 times the median of the absolute energy and no larger peak within 1.5 ms were considered spikes. The parameters for spike detection were chosen according to previous work [55]. After spike detection, an encoding method of choice (see Section 2.7.1 and Fig. S2) which maps a spike train during the response window to a scalar value was applied for each electrode. The resulting values were averaged across all recording electrodes.

### 2.7 Information Encoding

In this work, four different encoding schemes (see Fig. S2) for spike trains of each experiment run were evaluated and compared in terms of performance.

#### 2.7.1 Neural Codes

Rate coding was defined as the total number of spikes within the response window. ISI encoding was also used. There, the output is interpreted by calculating the average distance between spikes. ISI is not defined if there are less than two spikes detected within the given response time window. In these cases, the ISI response was set to the duration of the considered response time window. Classical phase coding could not be used for the experiments of this work because it requires an intrinsic background oscillation of the neuronal culture as a reference [25]. As only a short temporal segment is checked for a response, such an interpretation is not feasible. An inherited oscillatory behavior was not present in the cultures. Hence, the length of one cycle of the oscillation was defined as the time window considered. Afterwards, the average phase difference of all recorded spikes was calculated. In the case of zero spikes, the phase was set to 2*π−δ* with *δ* being the sampling interval. This ensures consistency with the handling of edge cases for other temporal codes. Another encoding type that is especially used with spiking neural networks is TTFS [56]. There, the delay between the stimulation and the first occurring spike was measured and all remaining activity was discarded. If no spiking occurred within a given time window, TTFS was set to the length of the analyzed time window. All four encoding schemes were computed for each electrode and subsequently averaged.

#### 2.7.2 Channel Capacity Estimation

To assess the performance of the different encoding schemes, notions from information theory were used. A stimulus at time *t* was modeled as a discrete random variable *X*_*t*_ and the neural activity induced by the stimulus was encoded to a scalar value by one of the introduced neural codes as the discrete random variable *Y*_*t*_. The probabilistic mapping from the stimulus to *Y*_*t*_ is called a channel. It was postulated that a good encoding contains most of the information of the input stimulus. The change of the entropy *H*(·), a measure of uncertainty, of *X*_*t*_ given the response *Y*_*t*_ was investigated. This is known as the mutual information:

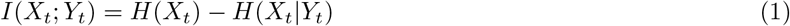

On one hand, it is beneficial to effectively use a large portion of the input alphabet. On the other hand, a symbol of the alphabet must be reliably transferred to the output. This optimization problem described by the mutual information has a theoretical limit of the capacity *C*:

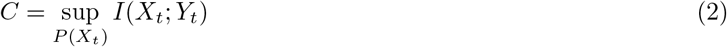

The capacity has the advantage over the mutual information as it does not assume any prior on *P* (*X*) and has the power to discard stimuli that only result in a highly variable output. For a discrete memoryless channel, it can be approximated with the Blahut–Arimoto (BA) algorithm [57, 58].

Memory was assessed by testing whether a time series 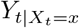 resulting from a pulse set is stationary with the augmented Dickey-Fuller test. Furthermore, significant active information storage (AIS) was evaluated with stateof-the-art estimators [59, 60, 61]. It was assumed that if there is no information gain of *Y*_*t*_ using *Y*_*t−*1_, there is also no information gain with *Y*_*t−i*_ *∀ i >* 1. Based on this assumption, AIS was estimated with a lag of one by computing *I*(*Y*_*t*_; *Y*_*t−*1_|*X*_*t*_ = *x, X*_*t−*1_ = *x*) using the Kraskov estimator [62]. The P-values across all stimulus parameters for both tests were combined using the weighted Stouffer’s Z Method. Time series with little activity (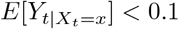 with the observable 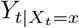 being rate encoded) were given a weight of one while time series with a larger average activity had a weight of ten. AIS was non-significant (*p >* 0.05) and the time series were stationary (*p <* 0.05), supporting the conclusion that the channel can be modeled as memoryless under the tested conditions. With this memoryless assumption, *t* is left out in the subsequent part.

To have a fully discrete system, it was necessary to partition an output observation *y ∈* ℝ into a region *b*_*i*_. For each experiment run and encoding separately, the edges of *n* bins were placed between the minimal and maximal value of the observed output *Y*. These edges were selected such that the frequency of observations was evenly distributed across all bins. An entry of the resulting transition matrix *T*, which fully describes the discrete channel, is defined as

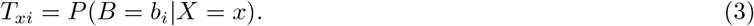

*P* (*X*) was estimated by maximizing the mutual information using *T* with the BA algorithm, which fully defines Ĥ (*X*). To approximate the conditional entropy term, the conditional distribution *P* (*Y X*) was estimated in a continuous fashion using a Gaussian kernel density estimation (KDE) with a bandwidth of 2.5% of the total span of the observed values *Y*. The estimate of the channel capacity becomes

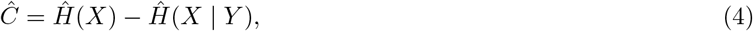

with

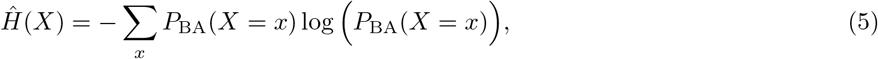

and

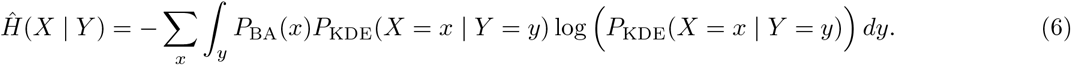

The number of bins *n* was chosen such that the bias was kept at a minimum while maintaining sufficient granularity (see Fig. S3).

## 3 Results

We report the successful long-term culturing of feedforward *in vitro* neural networks that can represent a nonlinear activation function (also referred to as transfer function), *i*.*e*., non-linear I/O relationship. By growing these networks inside custom-designed PDMS microstructures on top of CMOS HD-MEAs (see Fig. 1Bi), we can stimulate the network at precise presynaptic locations. We specifically chose the electrode at the start of the input microchannel for stimulation (see Fig. 1Bii), Bv), and Fig. 2A). This stimulation provides a means to reliably generate an action potential at the input, which then travels along the axons to the output neurons. The resulting input-output relationship constitutes a non-linear biological transfer function.

We demonstrate that these networks can perform basic computations, such as the NAND or OR gate. In addition, we explore how the type of information encoding in neurons influences its transmission, identifying the encoding method that maximizes information transmission and efficiency.

### 3.1 Information transmission is affected by choice of neural code

Since the mechanisms of neuronal communication remain unclear, it is yet ambiguous how to interpret extracellular recordings of neurons. To choose a good method of interpreting the output signal, we compared four neural coding schemes using the concept of channel capacity. We estimated the channel capacity using the approach described in Section 2.7.2 for response window sizes of up to 10 ms. The calculated averages across multiple networks of each neural coding scheme are shown in Fig. 3. We discarded networks that exhibit a smaller capacity than 1 bit per channel for any encoding from the analysis. In addition to the 10 ms response window, smaller variants were also evaluated. For both amplitude modulation (Fig. 3Ai) and frequency modulation experiments (Fig. 3Bi), TTFS coding outperformed the other encodings on average. As expected, most of the information is transferred during the very initial part of the response. As subsequent spikes are discarded, the noise resulting from more stochastic spiking at later stages is small, and the input can still be reliably transferred. Except for very short windows, rate coding performed slightly worse on average than TTFS coding for frequency modulation and significantly worse for amplitude modulation. There are two possible contributions to this observation. For longer windows, spikes triggered with lower probabilities are also taken into account. They lead to uncertainty in the transfer function. The occurrence of a spike for TTFS coding can alter the output value either continuously or, at a minimum, in discrete steps determined by the temporal sampling frequency of the recording system. For rate coding, such nuances are missing as the number of spikes recorded on an electrode is restricted to a small integer number due to the low cell count in the output node. This means that the accessible output range is rigid, and with limited output options, the transferred information is reduced.

**Fig. 3:**
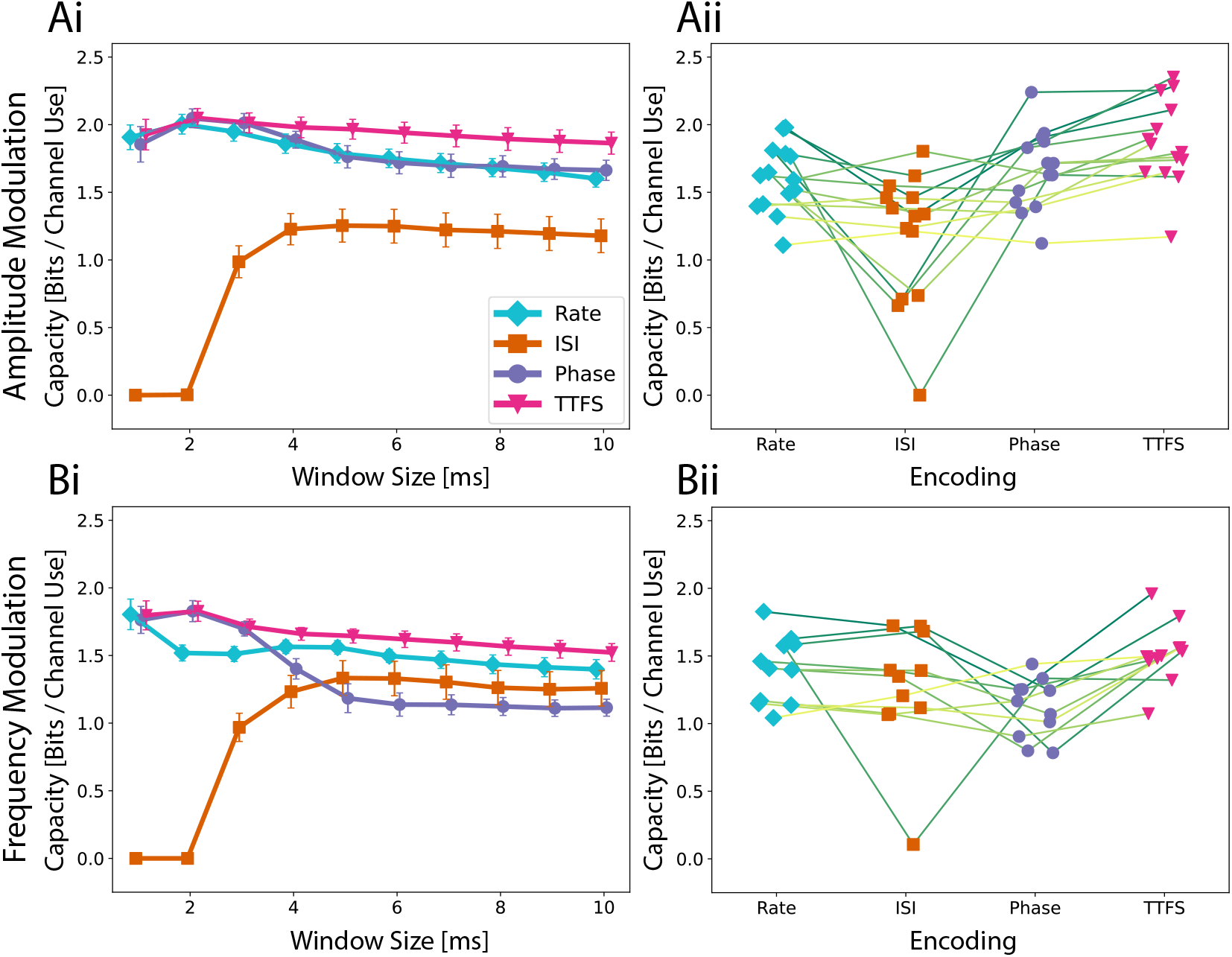
The Quality of a neural code is assessed with the estimation of the channel capacity. **(Ai)** Average capacity across fourteen networks as well as the standard error of mean for amplitude modulation. A higher capacity indicates that an encoding scheme is more expressive and reliable in transmission. The speed of the encoding is investigated by looking at multiple possible response window sizes up to 10 ms. **(Aii)** To investigate the performance differences on a network level, capacities for a response window of 10 ms are shown for each network. Encoding schemes corresponding to the same network are connected with a line. **(Bi)** Average capacity across ten networks for frequency modulation and **(Bii)** the analysis for a 10 ms time window on a network level.

Phase coding can be seen as an intermediate between TTFS and rate coding. While it does not suffer from the lack of expressiveness of rate coding, it can be more heavily affected by unreliable firing of neurons both in terms of spike occurrence and spike timing. This would explain why phase coding performed the worst for frequency modulation using a longer response window and yet was almost identical to TTFS coding when only the initial response was considered. For frequency modulation, the output inherently has a higher noise level. The capacity of ISI coding naturally behaves differently. This metric is not properly defined for small response windows as there is not enough time for multiple spikes to occur. Interestingly, ISI coding is highly robust to spiking noise occurring in later parts. While it failed to outperform any of the other neural codes for amplitude modulation, the obtained capacity for frequency modulation was comparable.

Capacity is not only determined by the encoding scheme. It also depends on inherent network properties. We observe three aspects when comparing the performances on a network level for a window size of 10 ms as seen in Fig. 3Aii), Bii). Phase coding is at most as good as TTFS. This suggests that spike timings in later window parts are in general not stable enough to carry additional information as seen in the decaying capacity curve. ISI coding either performs similarly to rate coding or worse. When compared to phase coding, ISI has a similar capacity for frequency modulation but tends to score lower for amplitude modulation.

In summary, the analysis underscores the importance of the initial response window size in determining the capacity of different encoding schemes, with TTFS coding matching or outperforming others with its ability to capture nuanced timing information. Rate coding suffers from a rigid, low-resolution representation, while phase coding strikes a balance between expressiveness and susceptibility to noise. ISI coding, though inherently unsuitable for small windows, demonstrates robustness to late-stage spiking noise, particularly in frequency modulation. These findings highlight the dynamic relationship between coding strategies and network properties in maximizing information transfer. In addition, information transfer during a later part of the response is more challenging due to the inherent stochasticity of the neural network formation and behavior. However, constraining neural growth within PDMS microstructures helps mitigate complexity and reduce variability to some extent.

### 3.2 Biological neurons exhibit non-linear responses similar to activation functions used in machine learning

To investigate the transfer functions in our feedforward networks, which consist of two inputs converging into a single output, we conducted experiments using both voltage amplitude and frequency sweeps as described in Section 2.6.3. In the first experiment, biphasic rectangular voltage pulses were applied with amplitudes ranging from 100 to 800 mV in 100 mV increments (see Fig. 2Bi) at a frequency of 4 Hz. Each voltage level was delivered simultaneously and identically through two electrodes located in the input axon channel (see Fig. 2A), in a sequence of 120 pulses. Since a few pulses were not sent due to hardware issues and/or their response was not detected in the post-processing, we decided to keep a total of 110 pulses for the analysis to ensure an equal number of repetitions for every amplitude. Neural spikes occurring within 10 ms after each stimulus were detected and plotted to generate a post-stimulus raster plot (see Fig. 2Bii), where each dot represents a neural spike on a corresponding color-coded output electrode, as defined in Fig. 2A.

To minimize possible carryover effects from prior stimulation, the order of the eight same-amplitude pulse sets was randomized, and a 5-minute rest period was included between stimulation sets. In Fig. 2Bii, we observed a stable response per electrode, reflected in the consistent timing of spike occurrences post-stimulus. This is manifested in the observed distinct vertical “activity bands” represented in various colors depending on the electrode location. Bands with shorter latencies appear more defined and likely reflect direct action potential propagation along the input axon. In contrast, bands with longer latencies are broader and paler, indicating a lower spike frequency and greater variation in post-stimulus latency, suggesting these are likely spikes from output neurons activated via synapses between the input axon and the output neuron’s dendrites. Since these spikes are the result of synaptic transmission rather than direct propagation of an elicited action potential, the stochastic component is more pronounced. They also occur less frequently, as the probability of successful synaptic transfer is less than 1, and are also less temporally precise due to expected variations in synaptic delay times [63, 64]. It is furthermore important to note that electrodes in the middle of the output region that are located below the seeding well and therefore uncovered by the PDMS have worse signal-to-noise ratio (SNR) as opposed to the covered regions covered (see Fig.S4). This means that activity from neurons positioned in this region will most likely not be recorded by the system, which is visible in Fig. S4 with the whole middle region of the output not having a spike recorded within the first 5 ms upon stimulation (electrodes colored in pink).

Additionally, as can be seen in Fig. 2Bii, we observed that stimulation voltages below 400 mV failed to elicit responses reliably in the output region, as indicated by the absence of activity bands. The recorded spikes are most likely due to spontaneous network activity. In contrast, voltages of 400 mV or higher consistently generated clear and regular response patterns in the output. However, the voltage amplitude at which the network reliably reacts to the applied stimulus is not necessarily identical for all networks, which will be discussed in more detail later. The observations are in line with previous work [45, 51].

For each electrode, the time of the first spike occurrence within a 10 ms latency window is taken and averaged across the electrodes in the output region, providing a mean TTFS per electrode. This encoding method is shown in Fig. 4Ai, where the mean TTFS per electrode is shown as a function of stimulation voltage amplitude. The results for the remaining encodings can be found in Fig. S5 Ai-Aiii. We observe a sigmoid dependency of the TTFS on the increasing stimulation voltage with a broad response distribution for amplitudes at or above 400 mV. In Fig. 4Aii, we present sigmoid fits for various individual 2-in-1-out networks (dashed lines) and the fit calculated over all collected data points for the experiment in question (solid line). While networks show reproducibility in the overall behavior, they display variability in saturation values and around the transition region, which is attributable to several factors:

**Fig. 4:**
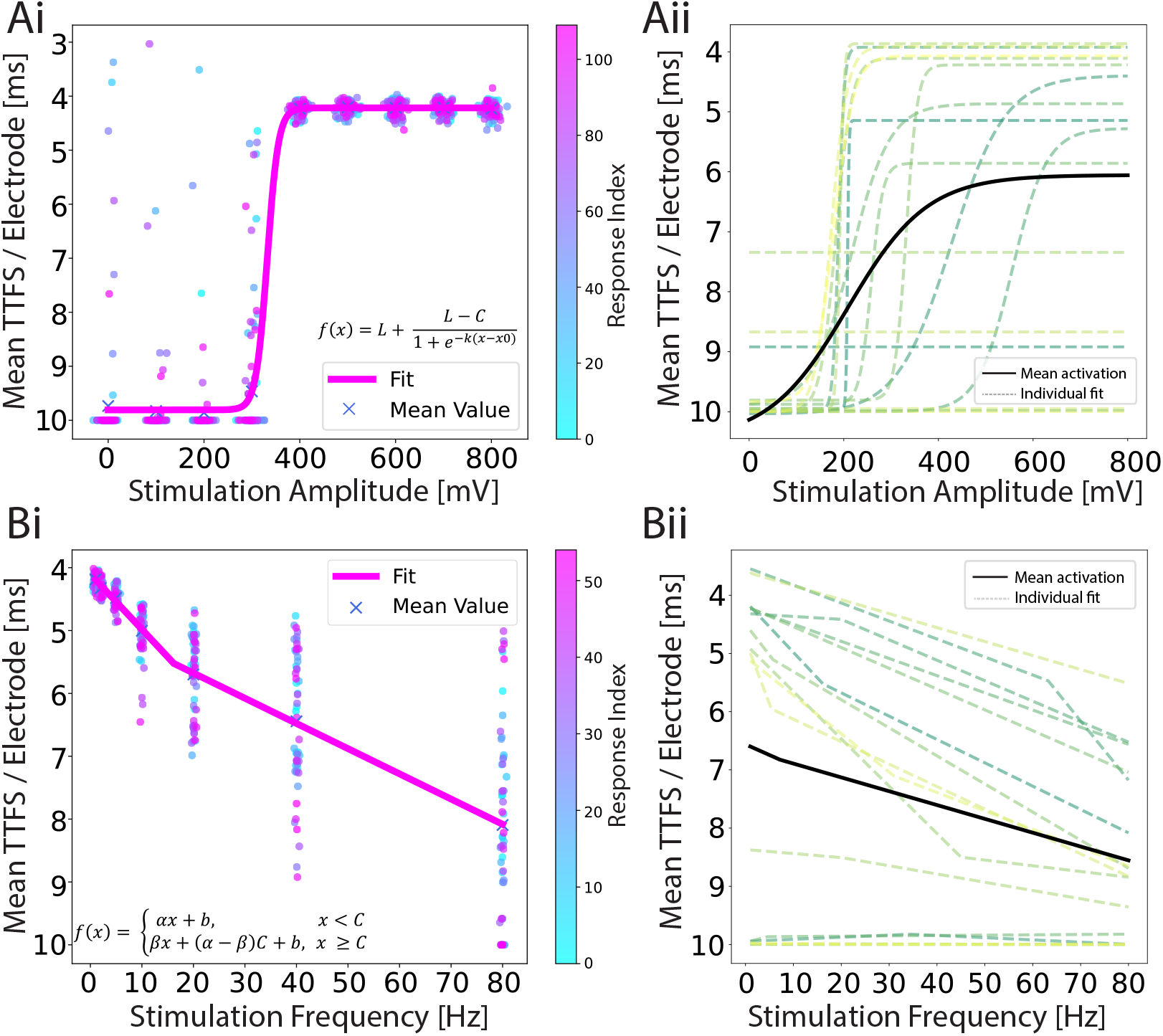
**(Ai)** An example of a response encoded as a TTFS for an amplitude-modulation experiment. A sigmoid function is fitted to a scatter plot of individual iteration responses. There are 110 responses for each amplitude and they are color-coded by time. **(Aii)** Individual sigmoid fits (dashed lines) for each network, and the mean sigmoid fit (black line) representing the average result from all experiments combined. Fit parameter values can be found in Table S3. **(Bi)** An example of a response encoded as a TTFS for a frequency-modulation experiment. Leaky ReLU is fitted to a scatter plot of individual responses. 55 responses are shown for each frequency and they are color-coded by time. **(Bii)** Individual leaky ReLU fits (dashed lines) for each network, and the mean leaky ReLU fit (black line) representing the average result from all experiments combined. In both experiments, the data is recorded from 17 networks and the curves are fitted using the non-linear least squares method. Fit parameter values can be found in Table S4. The y-axis is reverted in all plots.

The heterogeneity of neuronal networks introduces inherent variability in the input and output cluster characteristics. Due to the stochastic nature of cell seeding and potential cell loss during culture, cell cluster sizes differ, ranging from single cells to populations containing tens of cells (see Fig. S6). These spatial distribution and inter-connectivity differences can substantially influence neuronal activation patterns, particularly manifesting in variations of the sigmoid response curve’s plateau values. Additionally, axonal excitability is spatially dependent, with proximity to stimulation electrodes directly influencing action potential initiation. Networks with closer axonal adherence require lower stimulation voltages to trigger action potentials [65, 66], resulting in a variable input range for a transition region.

The overall sigmoid profile can be influenced by the choice of the stimulation electrode. In our approach, we recorded activity in the input channel region for 30 s and selected the electrode with the highest activity as the stimulation electrode. This criterion was intended to ensure that stimulation targeted an active axon, however, high activity upstream does not necessarily correlate with robust response elicitation downstream in the network.

In the second set of experiments, we applied a biphasic stimulation pulse at a fixed voltage amplitude but varied the frequency from 1 to 80 Hz, as shown in Fig. 2Ci. We picked 400 mV for the stimulation amplitude because this was the first amplitude above the threshold that we identified with the previous experiment. Seven frequencies were chosen: 1, 2, 5, 10, 20, 40 and 80 Hz. The number of iterations in the same time window for each frequency was in this case variable because it directly depends on the frequency. To ensure a fair comparison between the different parameters, the 1 Hz frequency pulse set was applied for twice the duration. For frequencies exceeding 2 Hz, a subset of responses was selected to maintain an equal number of samples for each parameter. Additionally, the time intervals between retained samples were standardized and maximized and only responses that resulted from stimulation on both input electrodes were kept. An example of a post-stimulus raster plot during this frequency sweep is presented in Fig. 2Cii. We observed higher output activity at lower stimulation frequencies (1–5 Hz), with a marked decrease in induced activity at higher frequencies. Additionally, the activity bands within the first 2 ms were more stable at lower frequencies.

These results suggest that high-frequency stimulation leads to activity depletion, a phenomenon previously noted in studies of deep brain stimulation at 130 Hz [67]. The frequencies that deplete activity (80–185 Hz) [68, 69] align with the higher frequencies used in our experiments. Other studies have demonstrated partial or complete blockage of neural activity, depending on the relationship between the stimulation frequency and the neuron’s intrinsic firing frequency [70].

The corresponding TTFS is given in Fig. 4Bi and shows an increase with increasing stimulation frequency. A leaky rectified linear unit (ReLU) is fitted to the data and shown in pink. The results for the remaining encodings can be found in Fig. S5Bi-Biii. In Fig. 4Bii we show the mean leaky ReLU of the 15 different 2-in-1-out networks. It is visible that for some networks stimulation evoked negligible activity (there were no spikes recorded within the 10 ms window on average), but for most networks, we observed the characteristic curve when encoding the output response in the form of TTFS.

In general, stimulation of 2-in-1-out neural networks shows reproducible behavior for both stimulation paradigms when stimuli with different amplitude or frequency are applied simultaneously at both input locations. Furthermore, amplitude and frequency sweep provide two distinct activation functions. The encoded output response is noisy due to intrinsic network activity as well as inevitable variations of the experimental parameters. For both input types, the response index shows a uniform distribution of response values over time (see Fig. 4 Ai and Bi), which means that we do not induce long-term modification in the network behavior with our stimulation paradigms. The insensitivity of the observed response to the randomization of the applied amplitude and frequency sequence also supports this claim.

### 3.3 Fundamental logic operations with biological neurons

In this section, we expand on the findings of Section 3.2 and now simultaneously apply either distinct voltage amplitudes or different frequencies to the chosen stimulation electrodes in the input channels. Additionally, we introduce an optional delay of 1.2 ms between the pulses of one iteration to investigate its effect on activation patterns (see Fig. S7). The latter pulse is considered the start of the response to mitigate potential issues of stimulation artifacts. The resulting activation patterns are illustrated in Fig. 5.

**Fig. 5:**
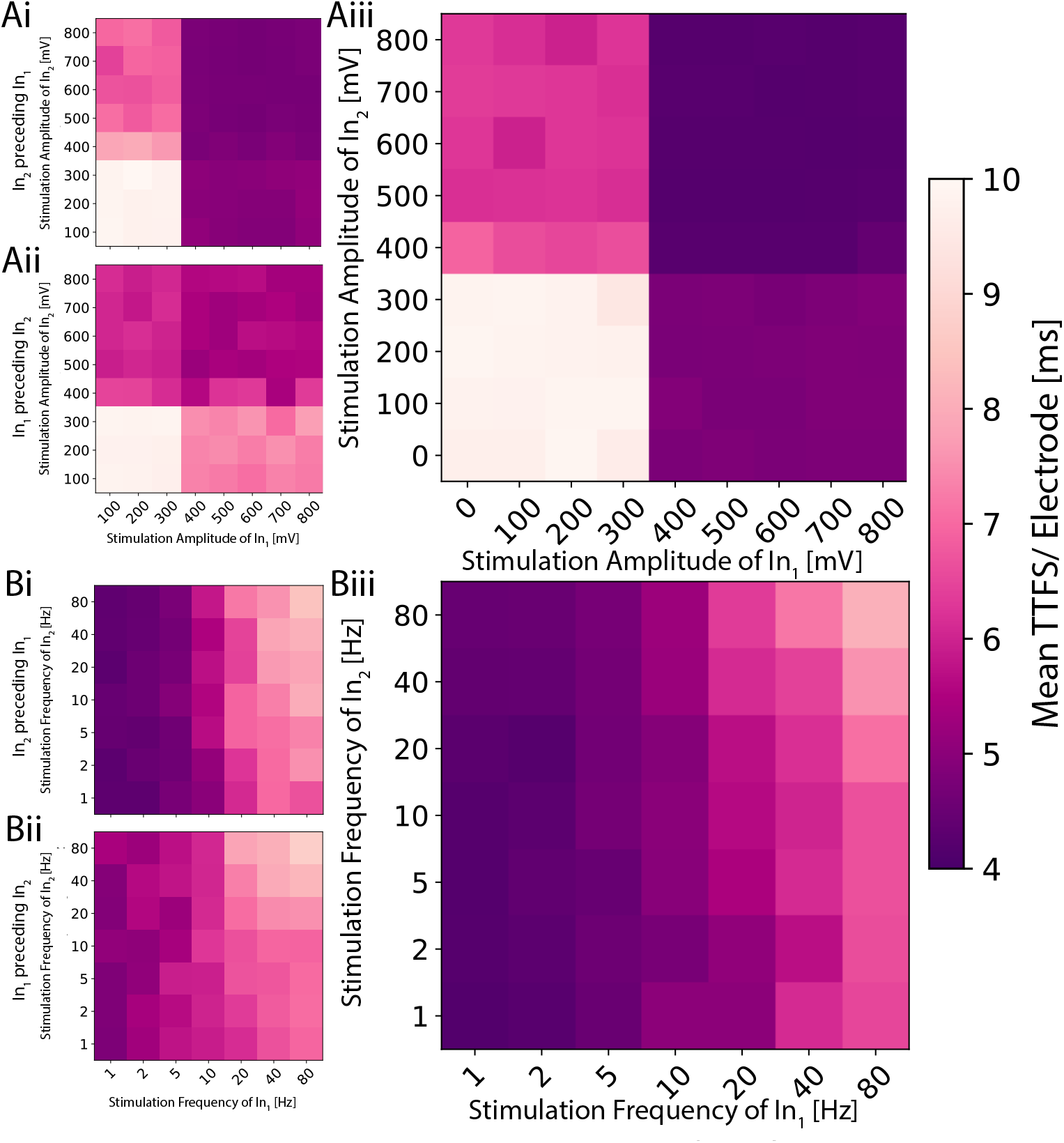
2D planes illustrating the activation function for distinct signals on the two inputs for a single network. **(Ai)** The amplitude is modulated on the two inputs independently, and the stimulation pulse sent to the second input precedes the one on the first by 1.2 ms. **(Aii)** The order is reversed. **(Aiii)** Both pulses are sent simultaneously. The resulting activation resembles a logic AND or OR gate depending on the applied threshold. **(Bi)** The frequency is modulated, and the periodic signal sent to the second input is shifted 1.2 ms earlier in time. **(Bii)** The shift is reversed. **(Biii)** Both signals are synchronized. The resulting activation exhibits low-pass behavior, analogous to a logic NOR or NAND gate depending on the used threshold.

As previously seen in Fig. 4Ai, voltages above a certain threshold elicited a slightly noisy yet consistent response for amplitude modulation when stimulating both electrodes with the same amplitude simultaneously. In this experiment, we investigated if and how the measured TTFS depends on which of the two pulses exceeds the threshold. We found four distinct regions within the activation function with high TTFS values in the bottom left quadrant, low values in the top right, and intermediate values in the remaining two quadrants (see Fig 5A). Analogies can be drawn to the operation of fundamental logic gates. In logic gates, multiple input bits are combined through logic operations to produce a single output bit. Similarly, if one maps the observed TTFS and the given input amplitudes to bits through thresholding, the network exhibits behavior akin to OR or AND gates. All tested neural codes showed an equivalent behavior as seen in Fig. S8. By applying a delay to one of the inputs, it is possible to shift the relative weighting between the two stimuli. The contribution of the earlier applied stimulus decreases as seen in Fig. 5Ai,Aii. This is in agreement with the observation seen in Fig. 2Bii that the initial response during the first few milliseconds gives a strong and stable band and becomes more sparse for later time points.

Frequency modulation exhibits a less evident pattern (see Fig. 5B) compared to amplitude modulation. For the TTFS encoding scheme, it shows a continuous low-pass filter in the mean. While still observable, this trend is less clear for the other three neural codes (Fig.S8). As previously seen in Fig. 4Bii, the response to higher frequencies spans a wide range of values. This means that it is not feasible to achieve a distinct response for each of those frequencies. However, the observation that responses are stable for low frequencies and that the average delay differs for high frequencies allows for the mapping of bits to frequencies in a way that can form logic NOR or NAND operations. For a delayed version of the input stimulus, the same observation is made as for the amplitude modulation. The latter input stimulus becomes the more dominant.

This behavior is consistent across different networks. For both modulations, one of the two stimuli has a larger impact compared to its counterpart. For the amplitude modulation, this means that the more influential stimulus gives rise to earlier spikes across the whole output region. For frequency modulation in general, high stimulation frequency results in low activity in the output. The input that has a weaker response for amplitude modulation will have severely less impact on the output for frequency modulation. This suggests that an input connection is responsible for a specific sub-region of the output. Overstimulated neurons of the first input node will no longer elicit a response in the neurons of the output node which can only be partially compensated by the neurons of the second input node. This would allow for more targeted control of a network’s output.

## 4 Discussion

This study demonstrates that the I/O relationship of small biological neuronal networks with controlled topology can be described as non-linear activation functions. These networks can perform basic Boolean operations, such as NAND and OR, which can be realized by network architectures consisting of two inputs and one output. This suggests that simple and compact neural architectures can achieve meaningful processing. These networks exhibit reproducible responses across different preparations and retain their computational capabilities over multiple weeks in culture, fulfilling essential requirements for scalability and potential applications in hybrid or bio-inspired artificial intelligence systems. Namely, the ability of biological neurons to perform non-linear activation and Boolean logic is analogous to how SNNs process and encode information using precise spike timing and thresholds. Observing and understanding these phenomena in biological networks provides a roadmap for improving SNNs, particularly in areas like robustness, adaptability, and real-time computation.

Furthermore, our analysis highlights the role of output encoding schemes in determining the quantity and quality of information transmitted through the system. Our work touches upon the limitations of rate coding for information transmission due to its reliance on temporal averaging, which can result in information loss. However, from the alternative spike-time output encoding schemes discussed here: TTFS, ISI, and phase coding, only TTFS outperforms it. This either implies that timing information is not captured in a sufficient way or that rate coding can capture quite some information even though it is the simplest encoding scheme. The latter point is also discussed by others [71], who argue that the rate vs. timing debate is not necessarily about which one is a relevant quantity, but rather whether rate can capture neural behavior accurately enough. This suggests that the selection of a specific coding scheme should align with the computational goals of the biocomputing system. These findings lay the groundwork for future research into more sophisticated and efficient biohybrid computational architectures.

There are long ways to go yet to fully realize the potential of biocomputing. Expanding the computational complexity of these networks is necessary to push their capabilities beyond basic operations. An important next step is the introduction of learning mechanisms, *i*.*e*., plasticity, through the development of stimulation paradigms for short-term and long-term modulation of synaptic weights. That could shift the paradigm from passively accepting the inherent behavior of the network to actively harnessing its computational capabilities for solving specific problems. The ability to modify biological synaptic weights would enable direct testing of the learning rules that have been commonly used in artificial bio-inspired systems [72]. Additionally, enabling real-time online interaction with the system will enhance control and adaptability [73, 55]. Input encoding schemes also require further exploration to optimize information transmission. Furthermore, to improve understanding and reduce variability, simplifying the network topology to study single-neuron I/O relationships could be important. Much of the current variability in our neural systems stems from inadequate control over network subpopulations and synaptic connectivity within and between neural nodes. Given that previous studies have demonstrated how network topology can influence the information processing capabilities of spiking neural networks [74, 75], addressing these methodological challenges to achieve comprehensive control and systematically correlate topology with computational characteristics is important for advancing hybrid intelligence and biocomputing system development. The adaptability of PDMS design platforms offers a promising experimental approach to empirically validate whether *in silico* proposed network topologies translate effectively into functional biological neural architectures.

To summarize, understanding how biological networks process information at a fundamental level enhances the plausibility and fidelity of bio-inspired AI systems. Incorporating these principles into AI models can improve their generalization, learning efficiency, and adaptability, aligning them more closely with human cognitive capabilities. The ability to execute Boolean logic is a foundational requirement for computation. If small biological networks can perform these tasks, it validates the potential of spiking and bio-inspired systems to handle more complex logical and algorithmic operations at scale. This study attempts to bridge the gap between biological insights and computational implementation, facilitating advancements in energy-efficient, scalable, and biologically plausible AI systems.

## Author Contributions

These authors contributed equally: JK & KV

JK: Conceptualization, Methodology, Software, Validation, Formal analysis, Investigation, Writing, Visualization

KV: Conceptualization, Methodology, Software, Validation, Formal analysis, Investigation, Writing, Visualization

HY: Validation, Formal Analysis, Investigation, Review & Editing

CV: Validation, Formal Analysis, Investigation, Review & Editing

SJI: Conceptualization, Methodology, Validation, Review & Editing

SW: Conceptualization, Methodology, Validation, Review & Editing

JV: Conceptualization, Review & Editing, Resources, Supervision, Project Administration, Funding Acquisition

## Acknowledgments

We thank Vaiva Vasiliauskaite for advice on statistical analysis and Benedikt Maurer for fruitful discussions regarding signal processing. We also thank Giulia Amos for creating an illustration of a CMOS chip.

