## Supplementary information for "Engineered biological neural networks as basic logic operators"

#### S1 Supplementary Figures

Figure S1 shows an example of a voltage map obtained upon attaching the PDMS microstructure to a chip substrate. We observe 10 independent networks that we can test on one chip. Due to the stochasticity of seeding cells in suspension and cell survival, nodes contain different numbers of cells.

Figure S2 depicts different output encoding schemes that were studied in Section 3.3. For each encoding, a 10 ms time window upon stimulation was considered. We studied four encodings: rate, time to first spike, inter-spike interval, and average phase.

Figure S3 shows that the bias of the capacity estimation method introduced is minimal for 128 bins.

Figure S4 shows the spatial distribution of TTFS for one network when it receives inputs of 5 Hz from one side and 10 Hz from the other side. The middle output region is where the PDMS well for cell seeding is positioned. While there are active neurons in this region, their signal is not picked up due to low SNR for open electrodes. The side output region is covered with PDMS that is 4 $\mu$m above the electrodes and improves the SNR significantly. Figure S4 shows the spatial distribution of TTFS for one network when it receives inputs of 5 Hz from one side and 10 Hz from the other side. The middle output region is where the PDMS well for cell seeding is positioned. While there are active neurons in this region, their signal is not picked up due to low SNR for open electrodes. The side output region is covered with PDMS that is 4 $\mu$m above the electrodes and improves the SNR significantly.

Figure S5 shows data points and sigmoid fit for the remaining three encodings: ISI, phase, and rate.

Figure S6 illustrates an example of hippocampal neurons cultured within the 2-in-1-out PDMS microstructures integrated on a CMOS chip at day in vitro 21.

In Figure S7 we depict stimulation delay paradigm. The delay between the two stimuli is varied between -1.2 and 1.2 ms.

Figure S8 shows the 2D planes of the activation function resulting from impaired stimulation for all four neural codes.

Figure S9 shows the results of the impaired stimulation experiments for additional networks.

Figure S10 shows the results of the impaired stimulation experiments for additional networks.

Laboratory of Biosensors and Bioelectronics (LBB), Institute for Biomedical Engineering, ETH Zurich, Zurich, Switzerland
* Authors have equal contribution.
‡ Corresponding author

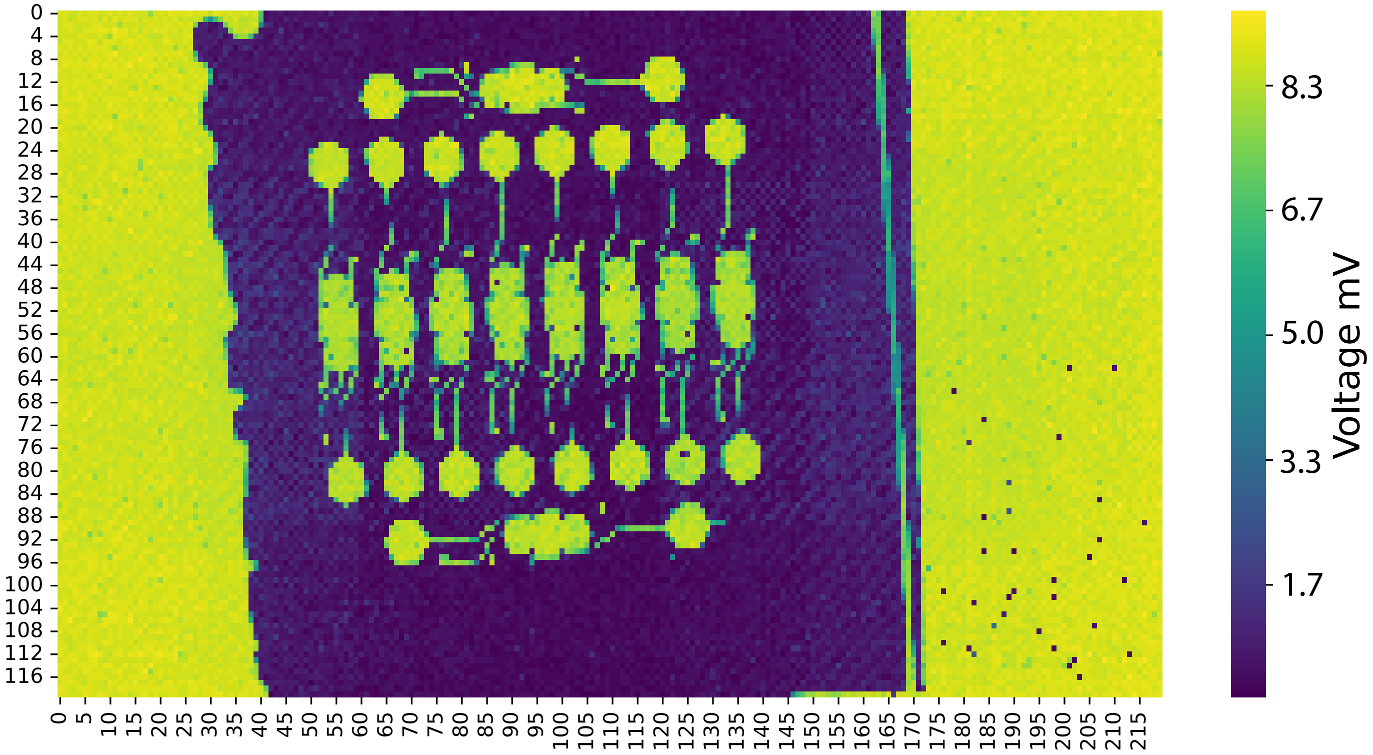


**Fig. S1**: An example of a voltage map obtained after microstructure adhesion to the chip. The difference in impedance between electrodes that are covered in PDMS and that are free enables us to identify the independent networks.

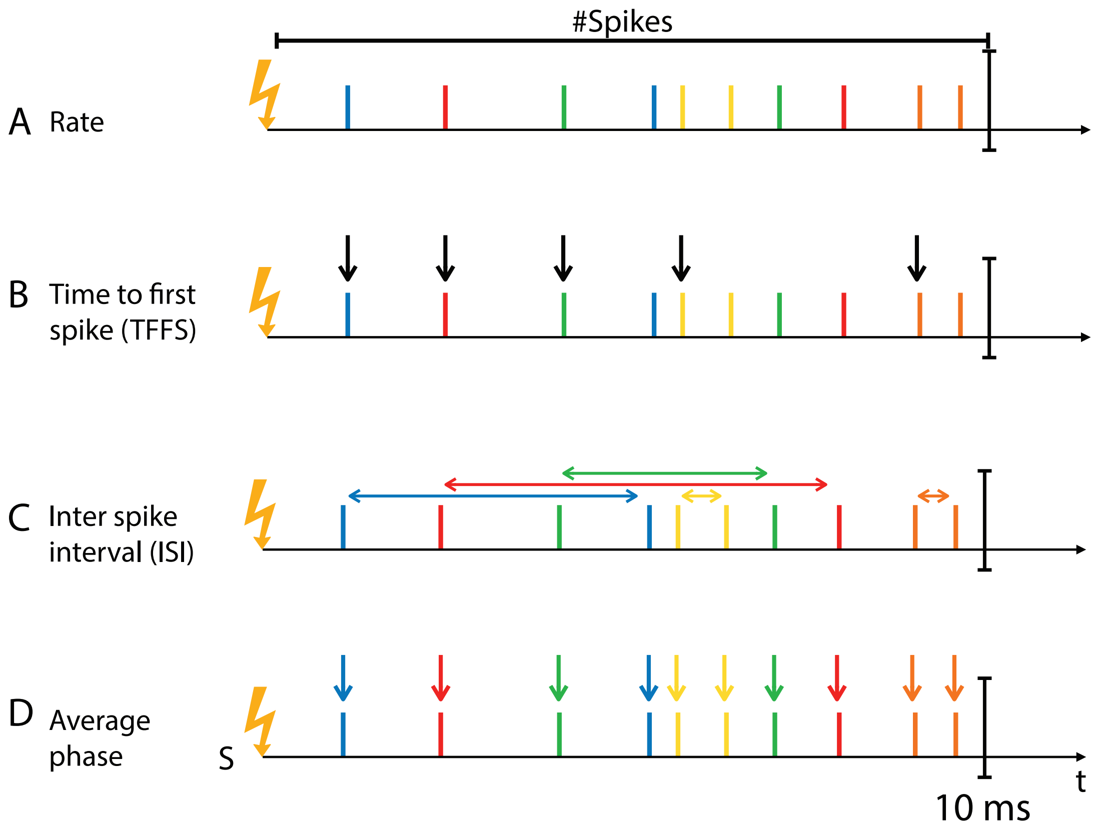


**Fig. S2**: Different output encodings for the defined 10 ms post-stimulus latency window. **(A)** Rate encoding includes counting the average spike per electrode. **(B)** Time to first spike corresponds to the latency of the first spike recorded on each electrode. **(C)** Inter-spike interval calculates the time difference between two consecutive spikes on each electrode. **(D)** Average phase coding assumes the internal reference oscillation.

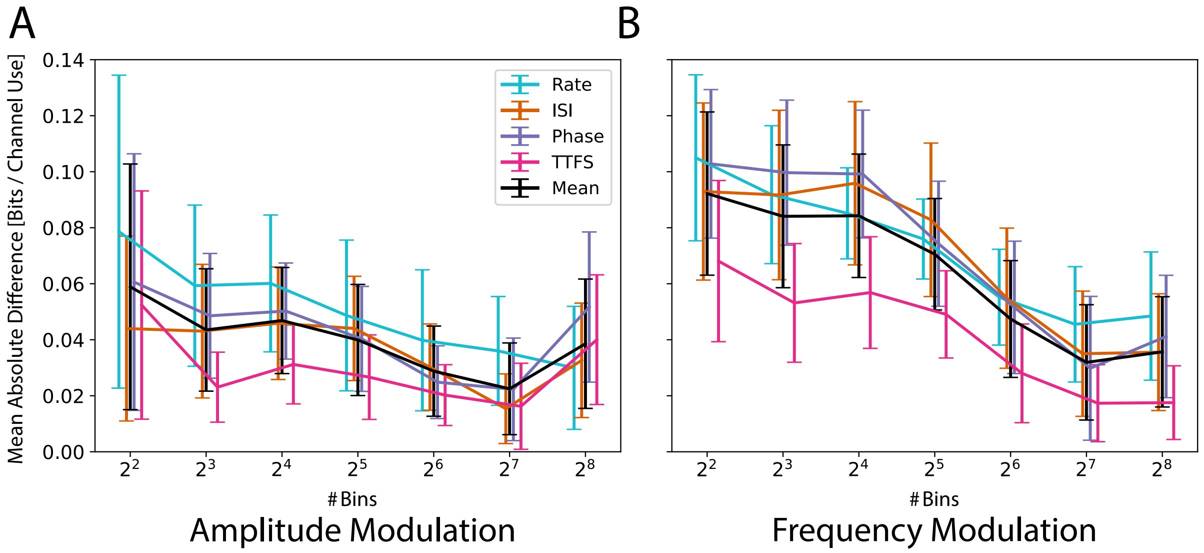


**Fig. S3**: Bias in capacity estimation for **(A)** amplitude modulation and **(B)** frequency modulation. The total dataset, composed of multiple pulse sets, was either kept intact or each pulse set was divided into two, three, or four non-overlapping subparts. For each division, separate datasets containing only the respective subparts were created, and capacity was estimated for each dataset configuration. The number of bins used for discretization was selected to minimize the difference in the average estimated capacity across all encoding schemes and dataset sizes.

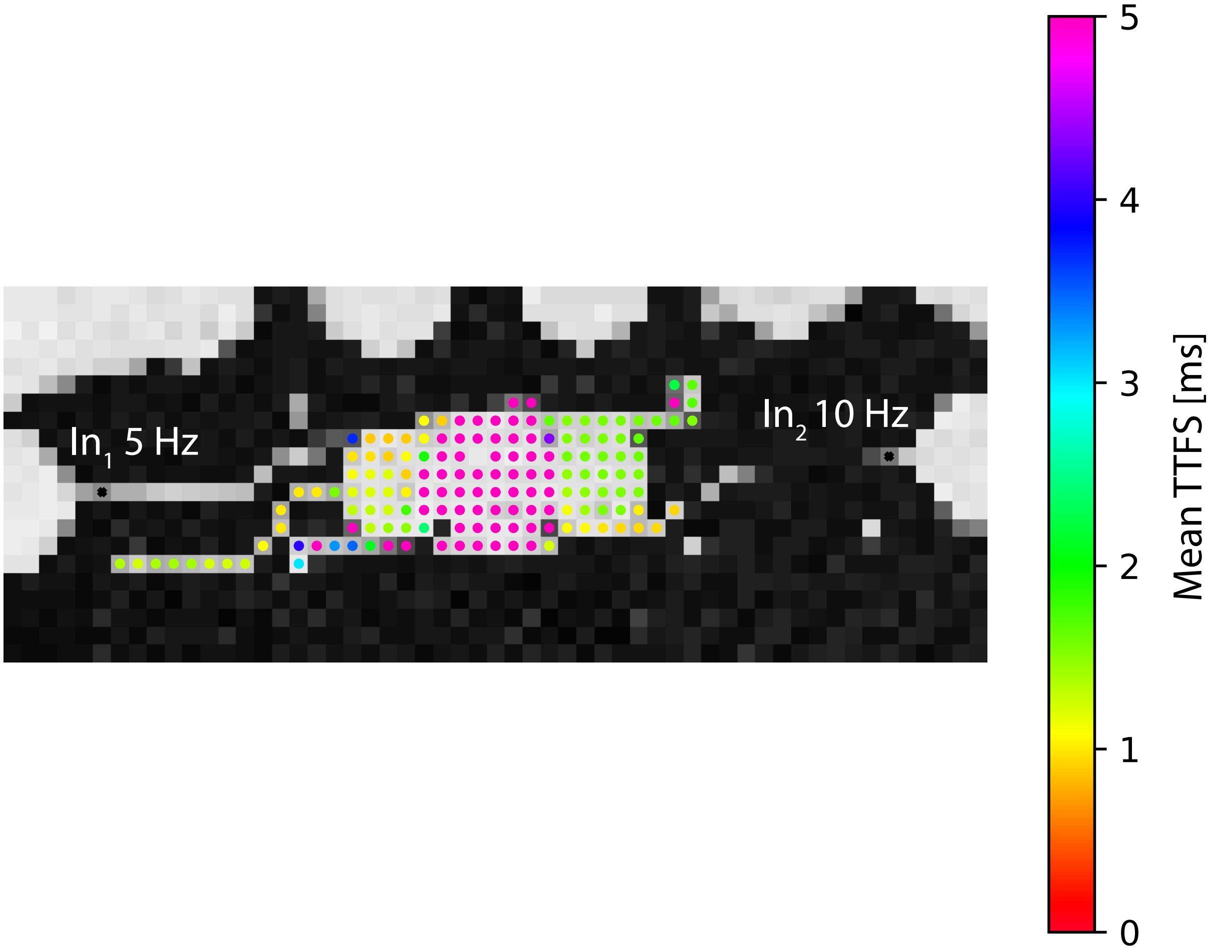


**Fig. S4**: Mean TTFS per electrode in the output region. The TTFS range is capped at 5 ms for improved visualization. The middle output region corresponds to the open microstructure, while the side regions are covered by PDMS positioned 2 $\mu$m above the electrodes. The network was stimulated with a frequency of 5 Hz from one side and 10 Hz from the other side, with a stimulation voltage amplitude of 400 mV in both cases.

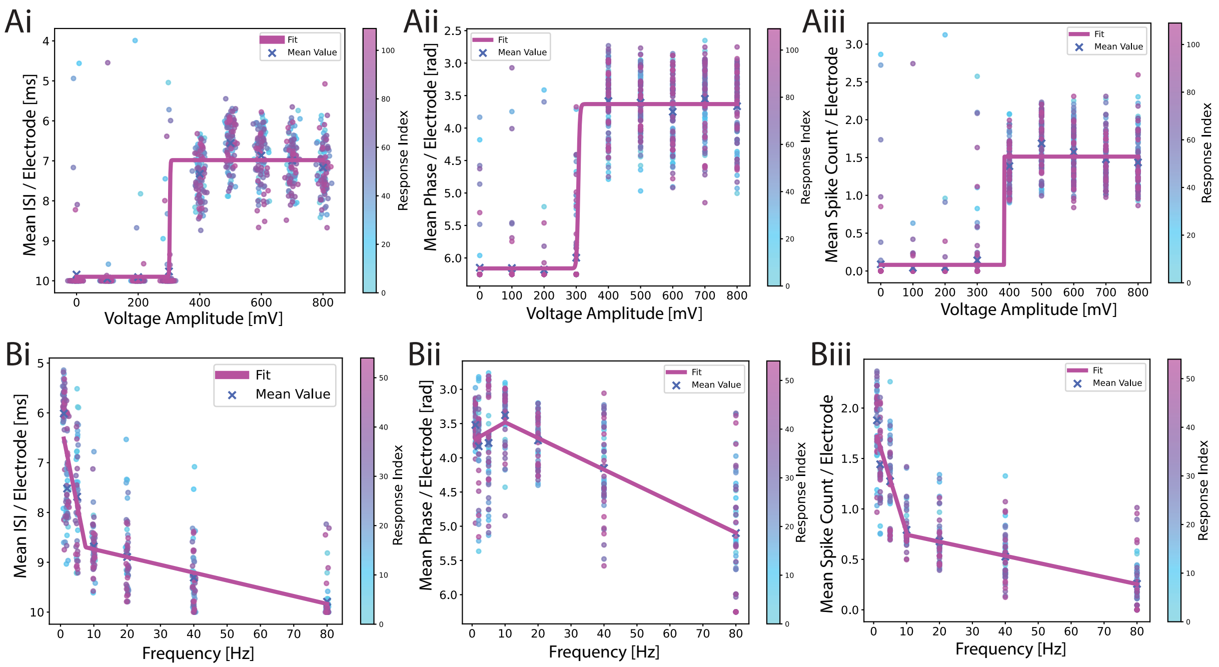


**Fig. S5**: **(A)** An example of a response for an amplitude-modulation experiment encoded as an average **(Ai)** ISI, **(Aii)** Phase and **(Aiii)** Rate output encoding. A sigmoid function is fitted to a scatter plot of individual iteration responses. There are 110 responses for each amplitude and they are color-coded by time. B) An example of a response for a frequency-modulation experiment encoded as an average **(Bi)** ISI, **(Bii)** Phase, and **(Biii)** Rate encoding. Leaky ReLU is fitted to a scatter plot of individual responses. 55 responses are shown for each frequency and they are color-coded by time.

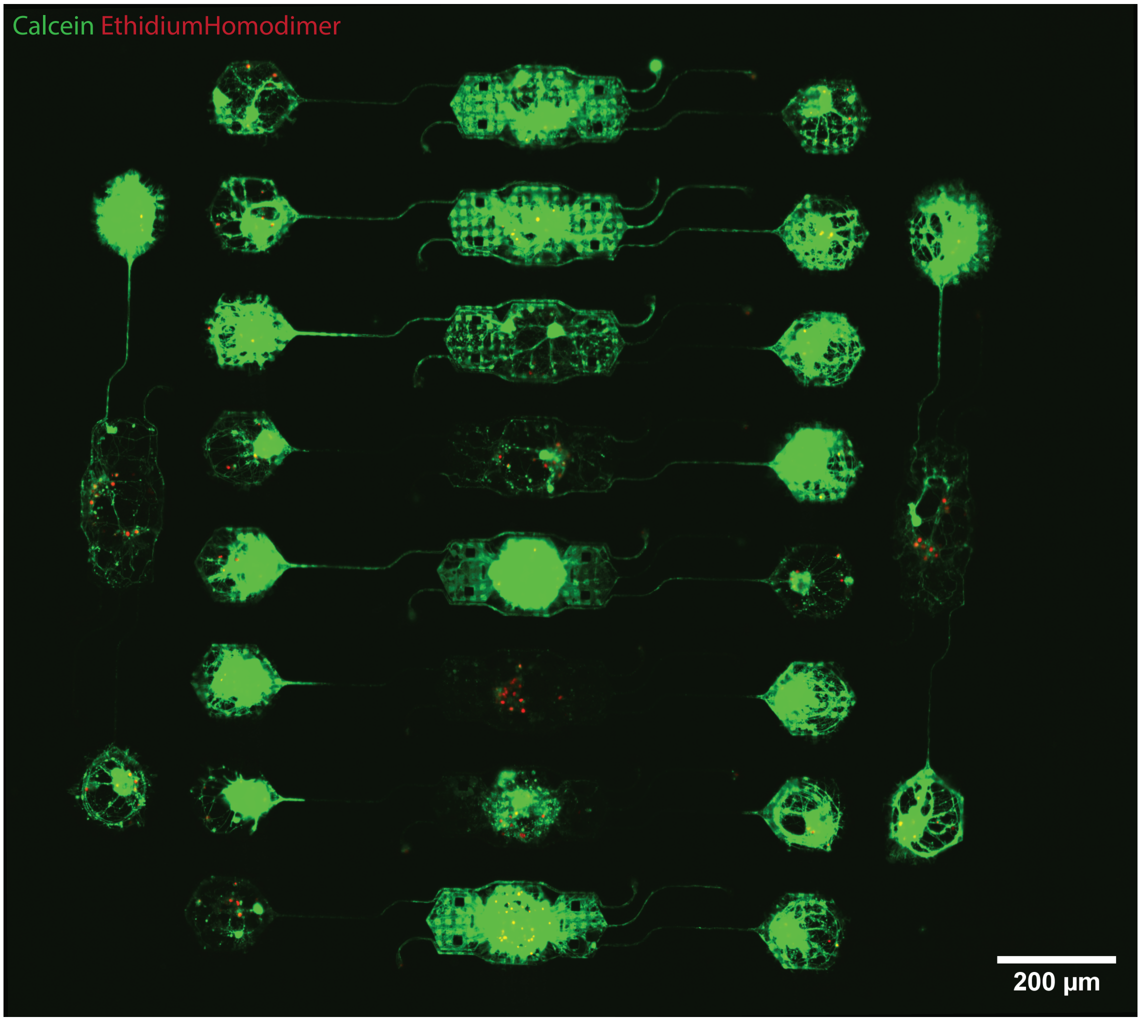


**Fig. S6**: A microscopy image of a single PDMS microstructure containing 10 independent networks on a CMOS chip. Hippocampal neurons at DIV 14 were stained with Calcein (green, indicating live cells) and Ethidium Homodimer (red, indicating dead cells).

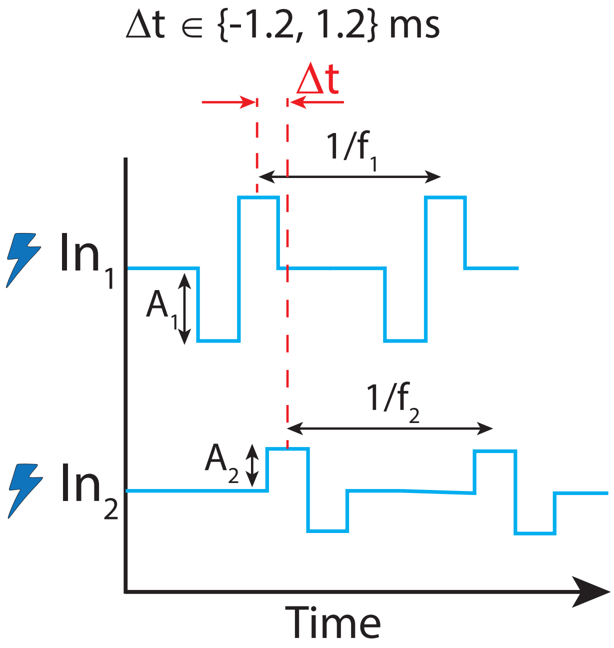


**Fig. S7**: An additional parameter added to both frequency and amplitude modulation experiments is a delay between stimuli applied to the two inputs. The delay has a fixed value of 1.2 ms with a varying sign.

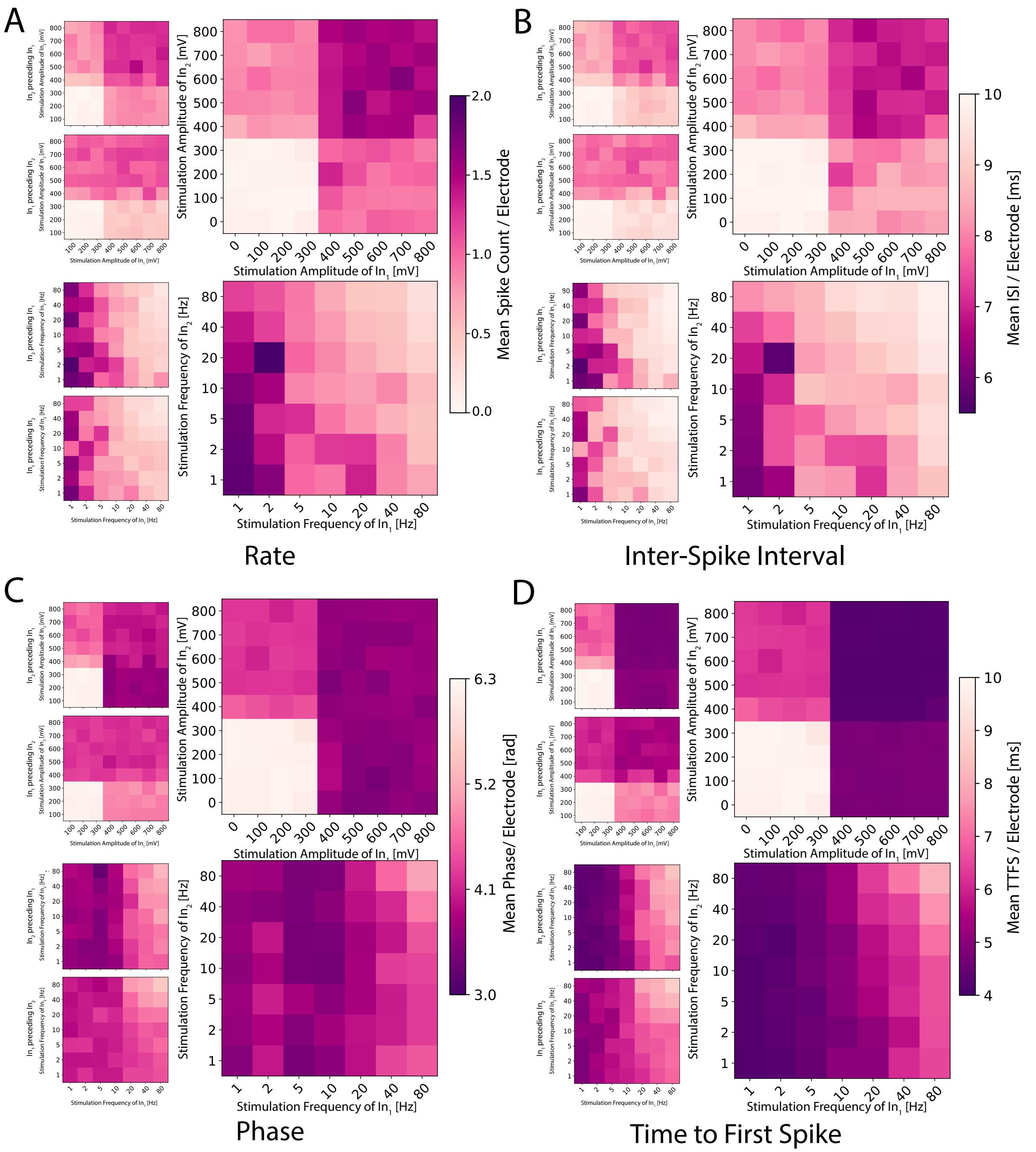


**Fig. S8**: 2D planes illustrating the activation functions for distinct signals on the two inputs for **(A)** rate coding, **(B)** ISI, **(C)** phase coding, and **(D)** TTFS.

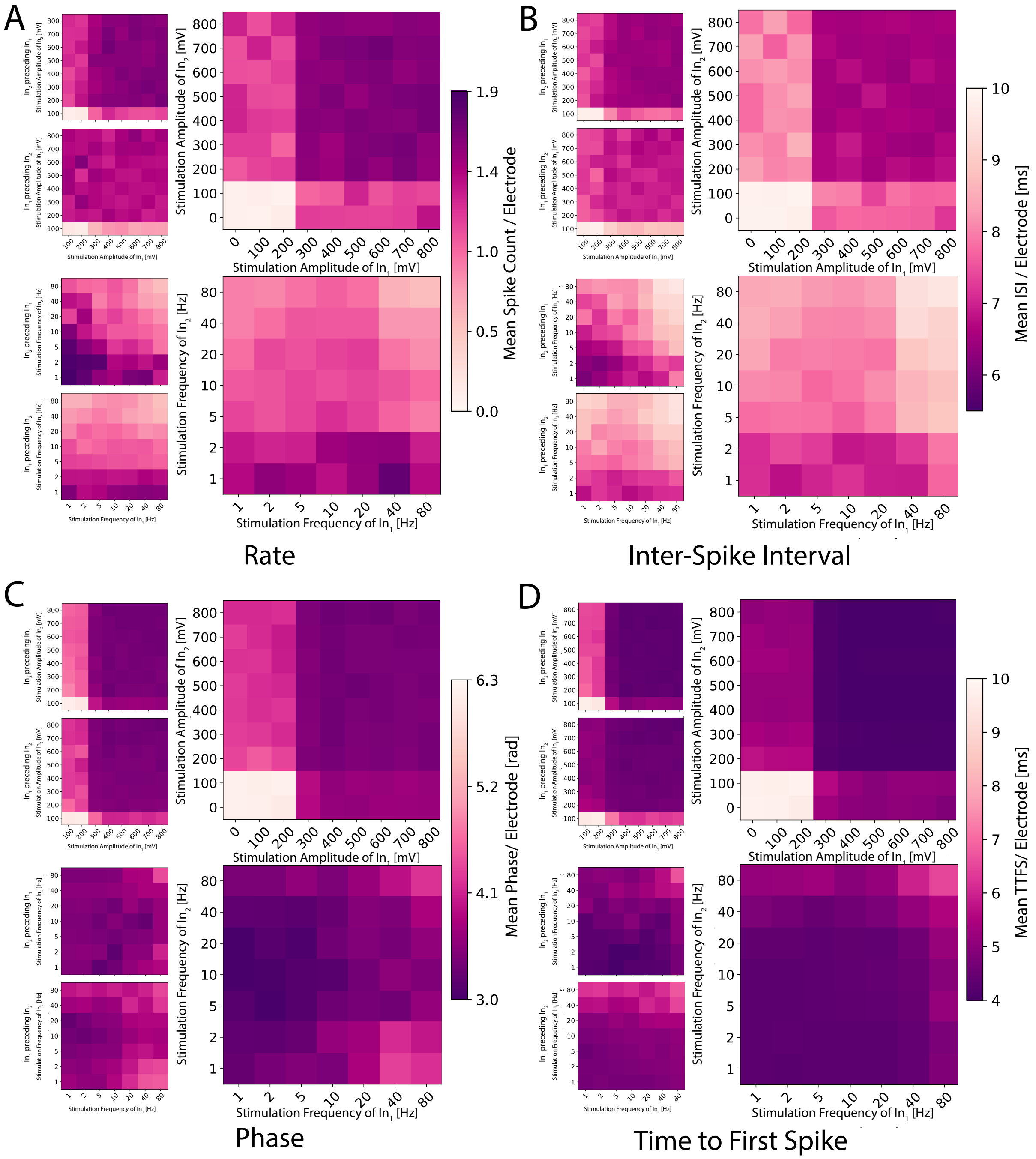


**Fig. S9**: Different network showing similar behavior to the one shown in Fig. S8.

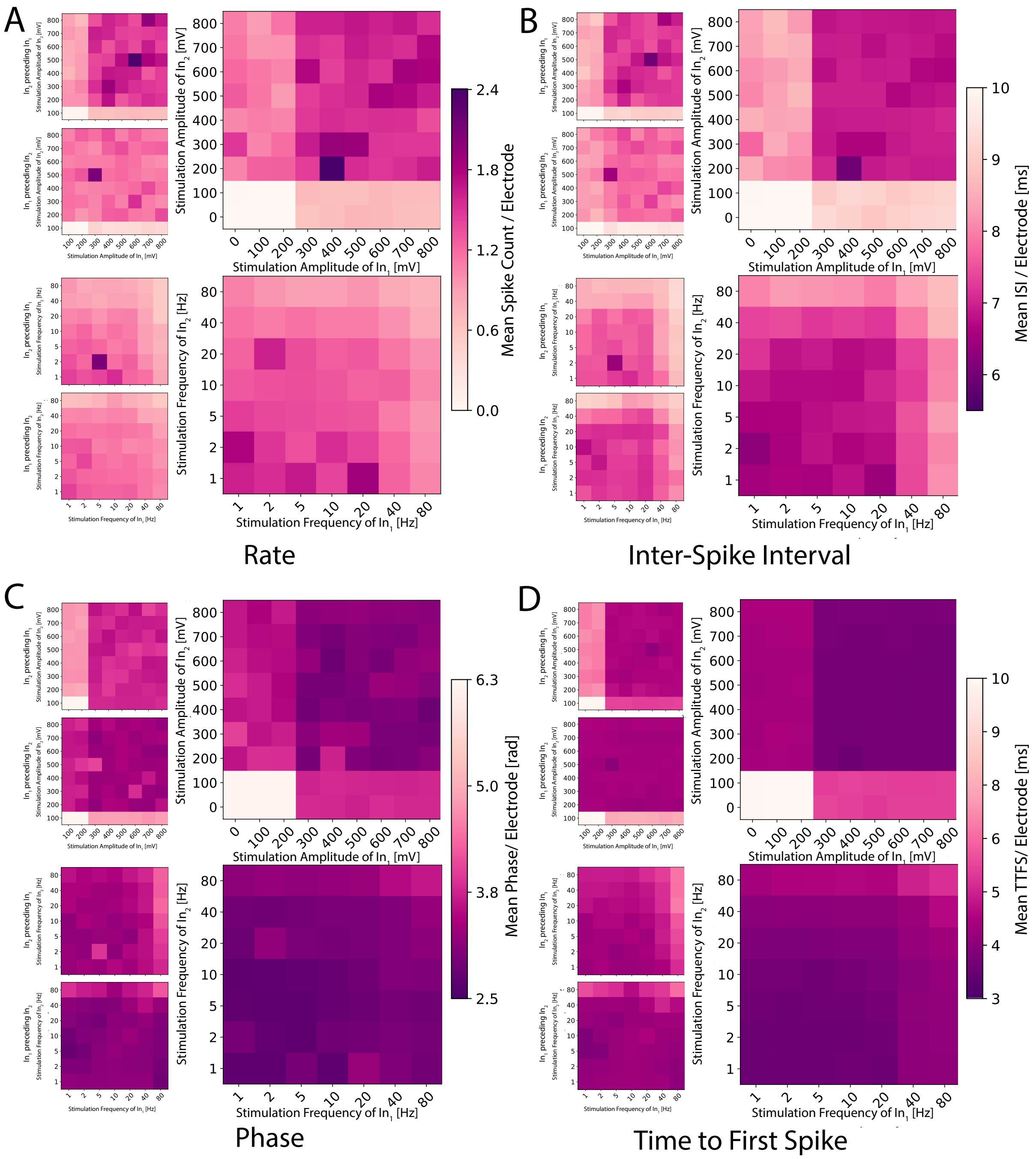


**Fig. S10**: Additional network showing similar behavior to the one shown in Fig. S8.

### S2 Supplementary Tables

| Pairwise Comparison | 1 ms | 2 ms | 3 ms | 4 ms | 5 ms | 6 ms | 7 ms | 8 ms | 9 ms | 10 ms |
|---|---|---|---|---|---|---|---|---|---|---|
| Rate vs ISI | **0.000** | **0.000** | **0.000** | **0.000** | **0.002** | **0.005** | **0.008** | **0.008** | **0.016** | **0.024** |
| Rate vs Phase | 0.903 | 0.125 | 0.639 | 0.808 | 0.626 | 0.542 | 0.670 | 0.952 | 0.808 | 0.268 |
| Rate vs TTFS | 0.429 | 0.087 | 0.650 | **0.005** | **0.002** | **0.002** | **0.004** | **0.003** | **0.002** | **0.001** |
| ISI vs Phase | **0.000** | **0.000** | **0.000** | **0.000** | **0.024** | **0.029** | **0.042** | **0.035** | **0.035** | **0.024** |
| ISI vs TTFS | **0.000** | **0.000** | **0.000** | **0.000** | **0.001** | **0.001** | **0.001** | **0.003** | **0.002** | **0.001** |
| Phase vs TTFS | 0.291 | 0.952 | 0.670 | **0.010** | **0.002** | **0.002** | **0.005** | **0.003** | **0.002** | **0.001** |

Table S1: P-values from the Wilcoxon test comparing capacity for amplitude-modulation between different neural codes across various response window lengths. Significant p-values are bolded.

| Pairwise Comparison | 1 ms | 2 ms | 3 ms | 4 ms | 5 ms | 6 ms | 7 ms | 8 ms | 9 ms | 10 ms |
|---|---|---|---|---|---|---|---|---|---|---|
| Rate vs ISI | **0.002** | **0.002** | **0.002** | **0.003** | **0.028** | 0.177 | 0.175 | 0.148 | 0.206 | 0.465 |
| Rate vs Phase | 0.438 | **0.006** | **0.015** | **0.021** | **0.004** | **0.014** | **0.027** | **0.027** | **0.037** | 0.064 |
| Rate vs TTFS | 0.700 | **0.004** | **0.029** | 0.122 | 0.177 | 0.081 | 0.101 | 0.148 | 0.185 | 0.152 |
| ISI vs Phase | **0.002** | **0.002** | **0.002** | 0.520 | 0.240 | 0.206 | 0.175 | 0.278 | 0.206 | 0.210 |
| ISI vs TTFS | **0.002** | **0.002** | **0.002** | **0.003** | **0.004** | **0.006** | **0.015** | **0.009** | **0.015** | **0.041** |
| Phase vs TTFS | 0.438 | 0.898 | 0.638 | **0.020** | **0.004** | **0.006** | **0.012** | **0.009** | **0.012** | **0.012** |

Table S2: P-values from the Wilcoxon test comparing capacity for frequency-modulation between different neural codes across various response window lengths. Significant p-values are bolded.

| Network # | $C$ | $L$ | $x_0$ | $k$ |
|---|---|---|---|---|
| 1 | 9.9 | 5.1 | 207.9 | 0.7 |
| 2 | 8.9 | 8.7 | -1175210.9 | -0.005 |
| 3 | 10.0 | 4.4 | 426.9 | 0.02 |
| 4 | 10.0 | 3.9 | 189.0 | 0.2 |
| 5 | 10.0 | 5.3 | 555.8 | 0.03 |
| 6 | 10.0 | 10.0 | 29890.6 | -210.6 |
| 7 | 9.9 | 4.9 | 215.9 | 0.02 |
| 8 | 9.9 | 5.9 | 258.0 | 0.08 |
| 9 | 9.8 | 4.2 | 332.0 | 0.09 |
| 10 | 10.0 | 4.1 | 183.7 | 0.06 |
| 11 | 7.3 | 6.6 | -2049914.7 | -0.0001 |
| 12 | 10.0 | 3.9 | 192.0 | 0.2 |
| 13 | 10.0 | 8.7 | 2827.8 | -16417.0 |
| 14 | 9.9 | 9.9 | -128375.4 | -706.3 |
| 15 | 10.0 | 3.9 | 173.4 | 0.06 |
| 16 | 10.0 | 10.0 | 10756.7 | -570.5 |
| 17 | 9.8 | 4.1 | 178.9 | 0.05 |
| Average fit | 10.5 | 6.1 | 208.9 | 0.01 |

Table S3: Parameters of the sigmoid curve fit obtained using *scipy.optimize.curve_fit*.

| Network # | $\alpha$ | $\beta$ | $C$ | $b$ |
|---|---|---|---|---|
| 1 | 7.3 | 0.001 | -9e7 | 10.0 |
| 2 | 16.2 | 0.09 | 0.04 | 4.1 |
| 3 | 34.2 | -0.004 | 0.004 | 10.0 |
| 4 | 63.2 | 0.03 | 0.1 | 3.5 |
| 5 | 20.0 | 0.005 | 0.04 | 4.3 |
| 6 | -10.3 | -0.05 | 0.03 | 3.4 |
| 7 | 4.6 | -0.02 | -0.0006 | 10.0 |
| 8 | -16.0 | 0.1 | 0.04 | 6.0 |
| 9 | 5.7 | 0.1 | 0.05 | 4.5 |
| 10 | 45.0 | 0.08 | 0.01 | 4.8 |
| 11 | 20.0 | 0.007 | 0.01 | 8.4 |
| 12 | 10.0 | -0.002 | 0.0002 | 10.0 |
| 13 | 1.3 | 0.09 | 0.02 | 3.5 |
| 14 | 5.4 | 0.2 | 0.04 | 4.8 |
| 15 | 31.0 | 0.07 | 0.03 | 5.1 |
| 16 | 2.0 | 0.02 | 3e6 | 10.0 |
| 17 | 21.4 | 0.001 | -2e-11 | 10.0 |
| Average fit | 7.2 | 0.04 | 0.02 | 6.6 |

Table S4: Parameters of the leaky ReLU curve fit obtained using *scipy.optimize.curve_fit*.

### S3 Artifact Removal

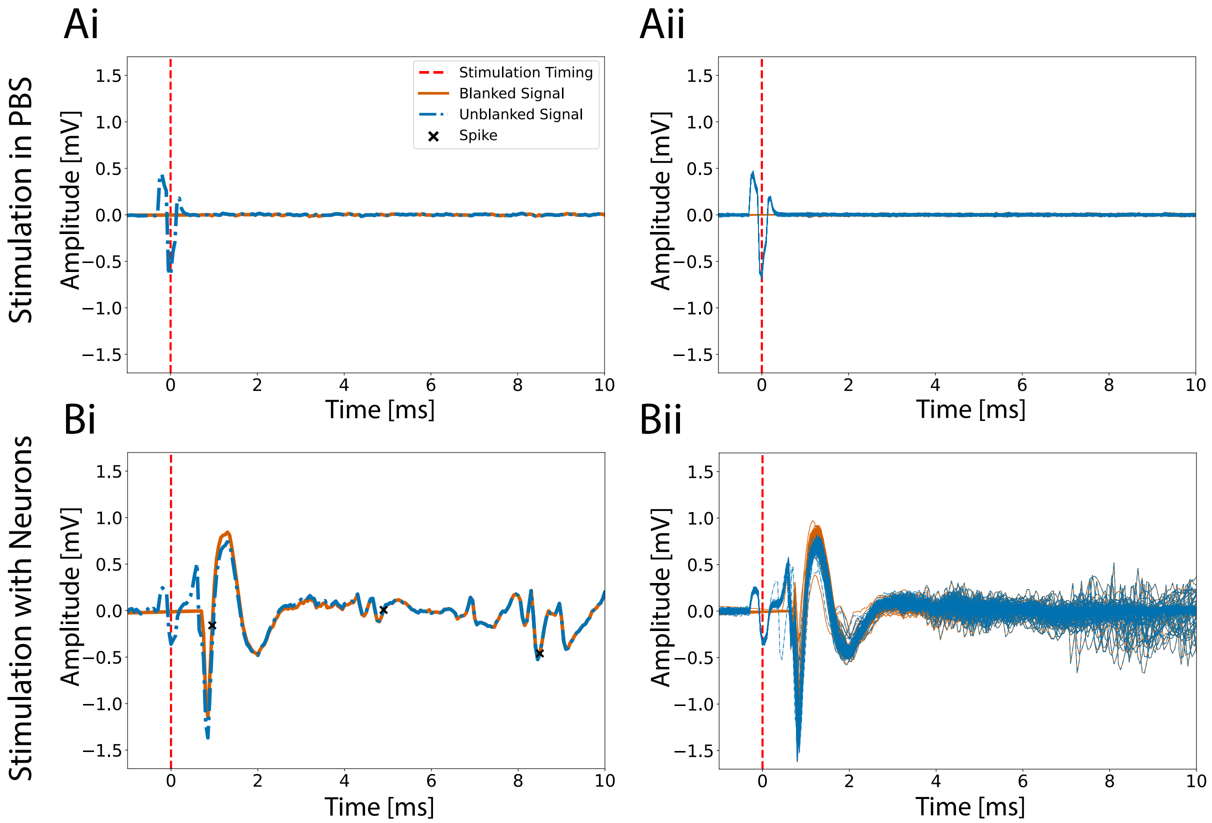


**Fig. S11**: Illustration of the artifact blanking. **(Ai)** Example trace of a stimulation event on a chip with the 2-in-1-out microstructure filled with PBS. **(Aii)** Overlaid traces of multiple stimulation events, demonstrating the consistency in artifact duration. **(Bi)** Example trace of a stimulation event on a chip with neurons. **(Bii)** Overlaid traces of multiple stimulation events on a chip with neurons.

An artifact blanking procedure was implemented to mitigate stimulation artifacts in the recordings. Stimulation timings were identified by detecting peaks in the recorded signals of the stimulation electrodes. The signals were filtered using a second-order Butterworth high-pass filter with a cutoff frequency of 200 Hz. The timing was determined by detecting peaks in the absolute value of the signal that exceeded 1 mV. For each recording electrode, the signal was filtered with a second-order Butterworth high-pass filter with a cutoff frequency of 20 Hz filter to eliminate low-frequency fluctuations. Subsequently, a signal window ranging from -0.25 ms to 1.5 ms relative to each stimulation event was extracted.

The shape of the resulting artifact was found to be influenced by the stimulus parameters but remained relatively consistent when these parameters were held constant. Therefore, for amplitude modulation, the artifact duration was extracted for each distinct pulse set individually. In the case of frequency modulation, artifact shapes were distinguished based on whether a pulse was sent on one or both input electrodes and the artifact duration was extracted accordingly.

Artifact duration was determined by computing the median of the extracted traces for each recording electrode, thereby obtaining a representative artifact shape while minimizing the impact of outliers. The time from the initial onset of the artifact to its return to the baseline, characterized by up to three peaks of alternating signs for recording electrodes sufficiently far away from the stimulation location, was measured. To further mitigate issues arising from stimulation-induced immediate shifts in the signal baseline, the blanking duration was extended by an additional 2 ms in the negative time. This duration was then removed from the unfiltered signal for each recording electrode around each stimulation timing, and the resulting gaps were filled using linear interpolation. An example of a blanked signal is illustrated in Fig. S11.
